# Neural activity profiles reveal overlapping, intermingled subpopulations spanning area borders in mouse sensorimotor cortex

**DOI:** 10.1101/2025.10.02.680044

**Authors:** Sohrab Salimian, Harrison A. Grier, Matthew T. Kaufman

**Author notes:** Equal contributions (Sohrab Salimian, Harrison A. Grier). **Data availability:** Data will be made available on FigShare upon publication. **Code availability:** Code used to create the figures is available on GitHub: https://github.com/kaufmanlab/SGK25-public.

## Abstract

Cortical control of movement is a distributed computation spanning multiple densely-interconnected regions. Although we have rich anatomical atlases and a coarse understanding of how function maps to areas and subregions, we lack a detailed account of how behaviorally-relevant activity is organized across the cortical sheet. Here, we trained head-fixed mice to perform a 15-target reach-to-grasp task while we performed cellular-resolution, two-photon calcium imaging across five regions of sensorimotor cortex (>39,000 layer 2/3 neurons). We characterized each neuron’s trial-averaged peri-event activity with interpretable metrics and mapped these response properties across areas, revealing large-scale spatial structure. Neuronal response profiles often shifted abruptly at anatomical borders: motor areas showed sharper tuning and more linear relationships with target location, whereas somatosensory areas displayed more heterogeneous response patterns. Neural response properties also differed according to somatotopic representation. Nonlinear dimensionality reduction of the neural feature matrix revealed that areas varied in their average response profiles, but that areas did not have well-separated feature distributions; instead, each area contained subpopulations. Neurons in each subpopulation had characteristic response profiles and were distributed across multiple cortical areas. The spatial distributions of the subpopulations overlapped, with neurons from different subpopulations salt-and-pepper intermingled in the overlap zones. Together, these results describe novel activity structure across sensorimotor cortex and identify several distinct but spatially-overlapping subpopulations with characteristic activity patterns during reach-to-grasp behavior.

## Introduction

Controlling movement is a critical function of the brain, and a sizable fraction of mammalian neocortex is involved in this task (Currie et al. 2022; Li et al. 2024; Muñoz-Castañeda et al. 2021). Moving in dynamic environments requires processes ranging from trajectory planning and pattern generation to error monitoring and feedback integration. Because of the importance of movement control, a wide variety of approaches have been used to study motor and somatosensory areas: from examining neural biophysics and connectivity, to determining neurons’ direct influence on movement, to relating activity of neurons in relevant subregions to behavior. However, to date the field has not systematically surveyed activity across sensorimotor cortex at single-cell resolution during behavior.

Anatomical mapping has formed the foundation of our understanding of the biological pieces of the brain’s puzzle, and has classically leveraged cytoarchitecture and projection tracing to describe how structure varies across the cortical sheet. These approaches support the view that cortex is divided into many areas that serve different functions, while remaining strongly interconnected (Brodmann 2005; Majka et al. 2021; Yeterian et al. 2012) especially in the mouse (Gămănuţ et al. 2018; Harris et al. 2019; Oh et al. 2014; Winnubst et al. 2019; Zingg et al. 2014). More recent genetic tools have complemented these methods with finer-grained insight into neuron-type distributions and connectivity, providing additional distinctions between parts of the brain (DeNardo et al. 2015; Muñoz-Castañeda et al. 2021; Wang et al. 2020) and revealing multiple parallel circuits within brain areas (Carmona et al. 2024; Park et al. 2022; Ueno et al. 2018). Subpopulations with distinct projection patterns and layer-specific connectivity suggest the existence of multiple functions coexisting within both somatosensory (Kerr et al. 2007) and motor areas (Carmona et al. 2024; Hausmann et al. 2022; Hira et al. 2013b; Muñoz-Castañeda et al. 2021; Zingg et al. 2014). Moreover, strong projections to subcortical, thalamic, and spinal targets originate from subregions of these cortical areas, some of which closely abide by anatomical borders determined by other methods (Carmona et al. 2024; Jeong et al. 2016) while others cross them (Hausmann et al. 2022; Ueno et al. 2018).

To relate activity to behavior more directly, other approaches have instead correlated neural activity with self-generated behavior. Large-scale recordings with simple behaviors have suggested that motor signals are widespread across cortex (Musall et al. 2019; Steinmetz et al. 2019; Stringer and Pachitariu 2019; Wang et al. 2023b), while studies with slightly more challenging behaviors have shown a greater proportion of motor signals in areas closer to the motor or sensory periphery (Li et al. 2024; Wang et al. 2023b). However, the details of what motor signals reside where have not been delineated. In an even more challenging motor task, neural populations in primary motor cortex (M1) and the primary somatosensory forelimb area yield similarly strong kinematic decoding of many tracked joint angles (Grier et al. 2026). These findings describe a distributed system, where signals are widely shared and where areas’ functional distinctions remain unresolved despite their sharp differences in connectivity with the periphery (Carmona et al. 2024; Ueno et al. 2018). A key question then remains: how are the activity patterns of individual cortical cells organized across sensorimotor cortex during behavior, and how does this relate to previous anatomical mapping?

To frame possible outcomes, consider that single neuron responses can vary along many dimensions. Cells could differ according to which movements or time periods they are recruited for (tuning), what movement parameters their activities reflect (encoding), or how their responses are structured across different movements (e.g., nonlinear encoding structure). Further, differences in these response properties across cells could be distributed over the cortical sheet in a variety of ways. Cells could form distinct “categories” or clusters that are spatially well-aligned to the boundaries of anatomically defined regions. Or, categories of neurons might span area boundaries in spatial footprints that do not relate obviously to area boundaries, and that either abut or overlap. At a fine-grained scale, cells with similar responses could be physically located near one another as in primate and feline visual cortex, or similarly-responsive neurons might be salt-and-pepper intermingled as seen in rodent visual cortex or in primate motor cortices during reaching behaviors.

Here, at single-cell resolution, we densely mapped single neuron activity across mouse sensorimotor cortex during a reach-to-grasp behavior to systematically capture the spatial distribution of tuning properties. We used two-photon calcium imaging to record from over 39,000 neurons as mice performed a 15-target reach-to-grasp task. We focused on five areas involved in this behavior: secondary motor cortex (M2), M1, and the primary sensory areas corresponding to the forelimb (S1-fl), hindlimb (S1-hl), and trunk (S1-tr). Our analyses emphasized the format of how neurons encoded information, rather than correlating activity with the behavior directly. Some tuning properties related to response onsets and peak timing closely followed anatomical borders. Other properties were more closely related to somatotopy, such as tuning sharpness, response duration, or the linearity of responses with respect to target location. Using bottom-up approaches, we were also able to identify subpopulations that shared response features, which were salt-and-pepper intermingled within anatomically-defined areas but had characteristic spatial footprints across cortex that spanned multiple areas. Together, these results clarify the organization of the response properties of neural populations at multiple scales, including identifying previously-unknown, overlapping and intermingled area-spanning subpopulations.

## Results

### Dense sampling of sensorimotor cortex reveals heterogeneous tuning

We implanted cranial windows covering sensorimotor cortex (Figure 1A-B) in six male mice expressing GCaMP8s in most cortical pyramidal cells (Zhang et al. 2023). Dense sampling of layer 2/3 neurons was achieved using two-photon calcium imaging across 98 fields of view (FOVs) spanning the forelimb, hindlimb, and trunk representations (Figure 1D). FOVs were aligned to the Allen Mouse Brain Common Coordinate Framework version 3 (Allen CCF) (Wang et al. 2020) using a combination of anatomical landmarks and widefield imaging with paw vibration mapping (Methods; Figure 1B). A total of 39,398 neurons were extracted across the five relevant regions in the sensorimotor cortex (Figure 1E). A larger number of neurons were imaged in M2 and M1 (31,453 neurons) than in somatosensory areas (7,945; Figure 1C).

**Figure 1.**
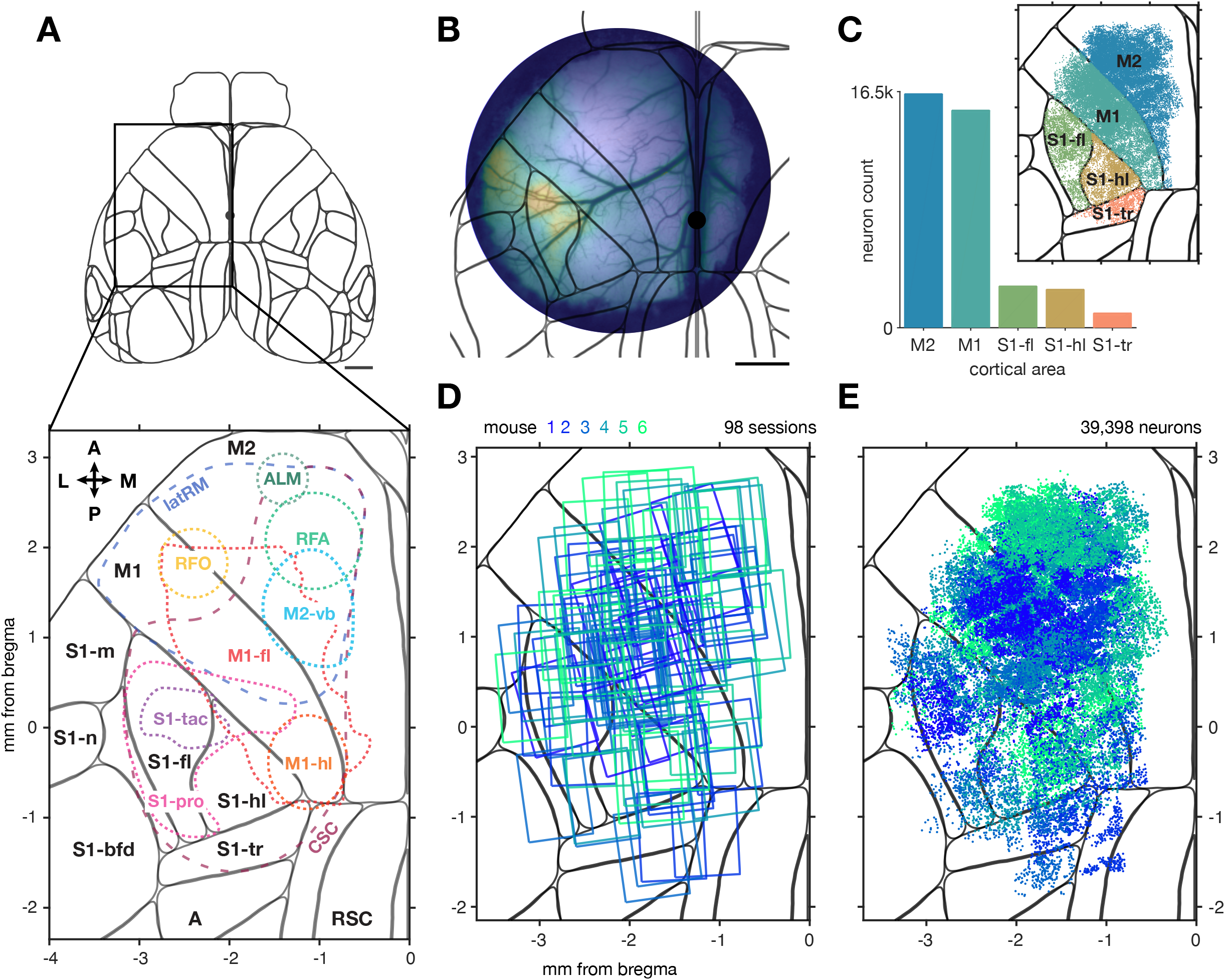
A dataset of densely sampled activity across mouse sensorimotor cortex. (A) *Top*, dorsal view of the Allen CCF, with the area investigated here indicated by a box. *Bottom*, summary of several previously-identified subareas of mouse sensorimotor cortex (see Methods for descriptions, alignment, and sources). Orientation legend: A, anterior, P, posterior, L, lateral, M, medial. (B) Widefield calcium imaging in a 5 mm cranial window superimposed on the Allen CCF to show location. Activation elicited by paw vibration (Methods). (C) Counts of neurons imaged for each area, with identities colored in inset. (D) Field of view locations of the 98 sessions of two-photon imaging collected in layer 2/3 for 6 mice (colors). (E) Locations of the 39,398 neurons extracted from all sessions. Mouse identity colored as in D.

#### Behavior

While neural activity was acquired the mice performed a trained, motorically-challenging, delayed reach-to-grasp-to-drink task (Methods). The task was a modified version of the Galiñanes and Huber (2023) multi-target reaching task. Mice were head-fixed and held two paw rests to initiate a trial (Figure 2A,B). A trial began with a spout moving into one of 15 positions, selected randomly on each trial from a concave grid arranged around the animal’s snout. After a variable delay period a non-target-specific auditory Go cue (“cue”) was played and a water droplet released, and the mouse could use its right paw to reach toward the target and retrieve the water reward. Stereo high speed camera data enabled markerless tracking (DeepLabCut; Mathis et al. (2018)) offline. Animals were successful at contacting the target on 97.1% of trials, and an average of 373 successful trials were collected per session (264-450; 39,229 successful trials total in the dataset).

**Figure 2.**
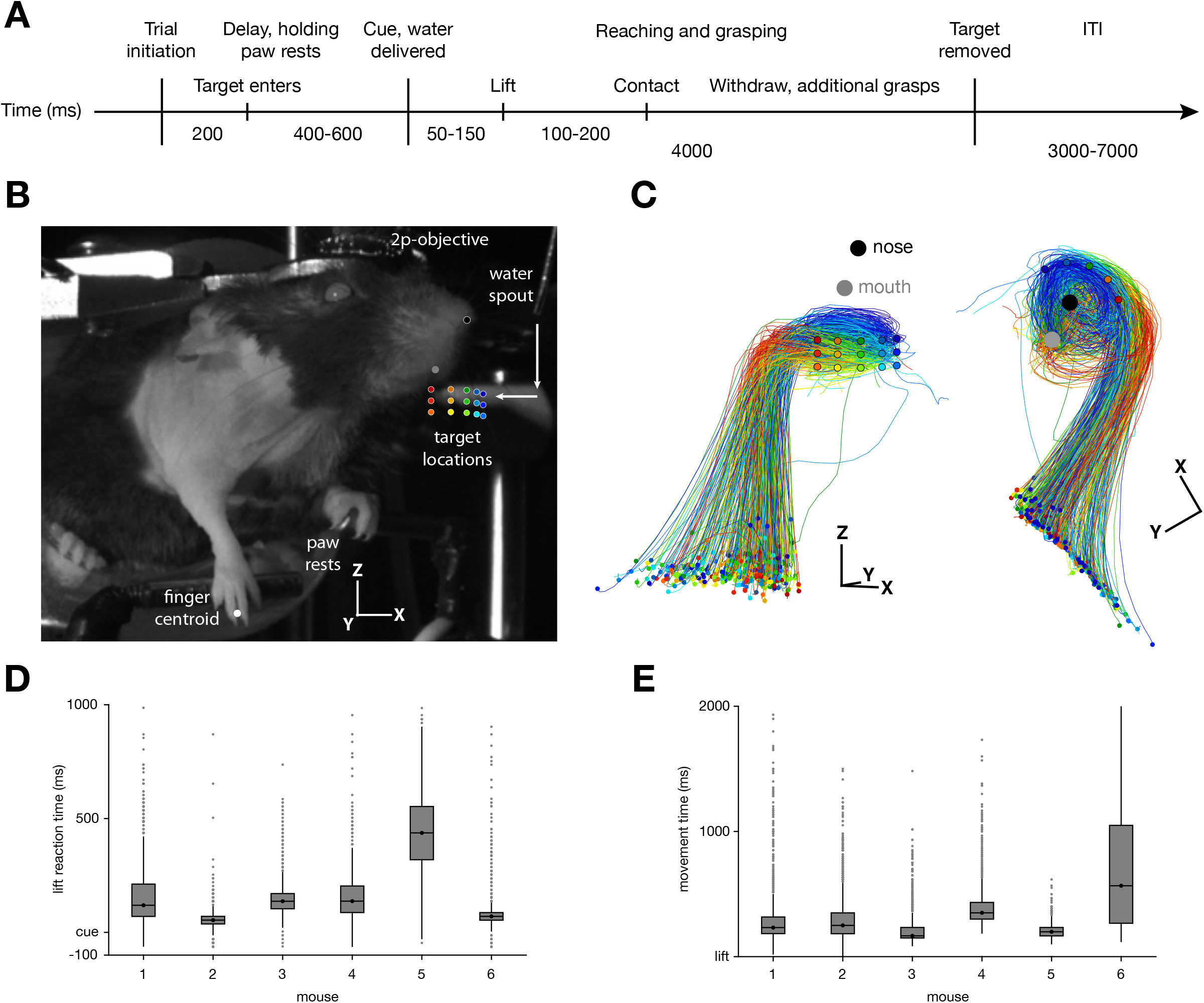
Mice performed delayed reach-to-grasp-to-drink movements to 15 distinct targets. (A) Task timeline. ITI, inter-trial interval. (B) Infrared image of the mouse during the inter-trial interval from one of two high-speed cameras. The 15 possible target locations illustrated with colored dots. (C) Finger centroid trajectories locked to lift onset from a frontal (left panel) and dorsal (right panel) view, illustrating kinematic separation by target. Target locations and average nose and mouth locations superimposed over centroid traces for clearer visualization. Coordinate axes show 3 mm scale bars and correspond to the axes in B. (D) Box plots of lift reaction times (RTs) for each mouse. Boxes show interquartile range (IQR) with a line for median; whiskers show 1.5 times the IQR; dots show outliers. Negative RTs indicate reaches before the Go cue, and were excluded from further analysis. (E) Time between lift and spout contact across mice. Three consecutive sessions per animal shown for both D and E.

As a simple characterization of the behavioral kinematics, single-trial paw centroid trajectories were plotted and colored by their corresponding target position (Figure. 2C and Figure 2-figure supplement 1A). Two relevant features were readily apparent. First, the paw centroid location was strongly target-specific near the spout, demonstrating that reaches were in fact targeted. Second, a looping trajectory near the spout was typically present, which corresponds with repeated reaches to collect all the water from the spout. This was due to a combination of our custom-shaped spouts, which were designed to cause the water to cling, and our use of touch sensors which encouraged strong contact with the spout itself (Methods). Although variability was present in both reaction time (Figure 2D) and movement time (Figure 2E), as is typical in reaching data, both intervals were generally short and indicated that animals were highly engaged in the task. To assess the extent to which mice aimed their reaches to each target, we computed the mean Euclidean distance across trials within target and between targets (Figure 2-figure supplement 1B). Reaches made to the same target were more similar to one another than were reaches to different targets, suggesting that the animals intentionally reached to each spout location rather than generically reaching out and searching for contact. We do not have data on how the mice localized the spout on each trial, but it could potentially have been by sniffing (Galiñanes et al. 2018), whisking, or detecting vibrations from the motors that positioned the target. A fuller characterization of how neural activity relates to the kinematic output will be detailed in forthcoming work.

#### Neural data visualization

To begin investigating the neural activity, we identified which neurons modulated their firing rates during the task. To do so, we computed a modulation score for all cells using a variant of the ZETA method (Methods; Montijn et al. (2021)), which asks whether the activity for a neuron changes over time consistently in relation to any given target. Modulated neurons were common and scattered across all five regions of interest in the sensorimotor cortex (Figure 3A). To quantify their spatial distribution, we computed the fraction of neurons that were modulated across the cortical sheet. The five areas of interest varied in the fraction of neurons that were modulated: M2 had 14%, M1 had 23%, S1-fl had 30%, S1-hl had 25%, and S1-tr had 27% (p < 10^−16^, Chi squared test for homogeneity, cue-aligned; between-area pairwise two proportion z-test in Figure 3-figure supplement 1B; Methods). Given that these areas have known spatial organization within them and structure was apparent by eye in the spatial scatterplot of modulated neurons (for example, the less modulated band along the M1/M2 border in Figure 3A), we then produced a high-resolution map of the fraction of neurons that were modulated by location (Figure 3B). Neurons in two zones were more often modulated than elsewhere: a wide strip along the border between S1 and M1, and an anteromedial patch of M2. The probability of a neuron being task-modulated was highest around the border between M1 and S1-fl, corresponding to the center of the forelimb portion of M1 and the proprioceptive region of S1-fl (Alonso et al. 2023). Probability of modulation was particularly low in a wide strip along the border between M1 and M2.

**Figure 3.**
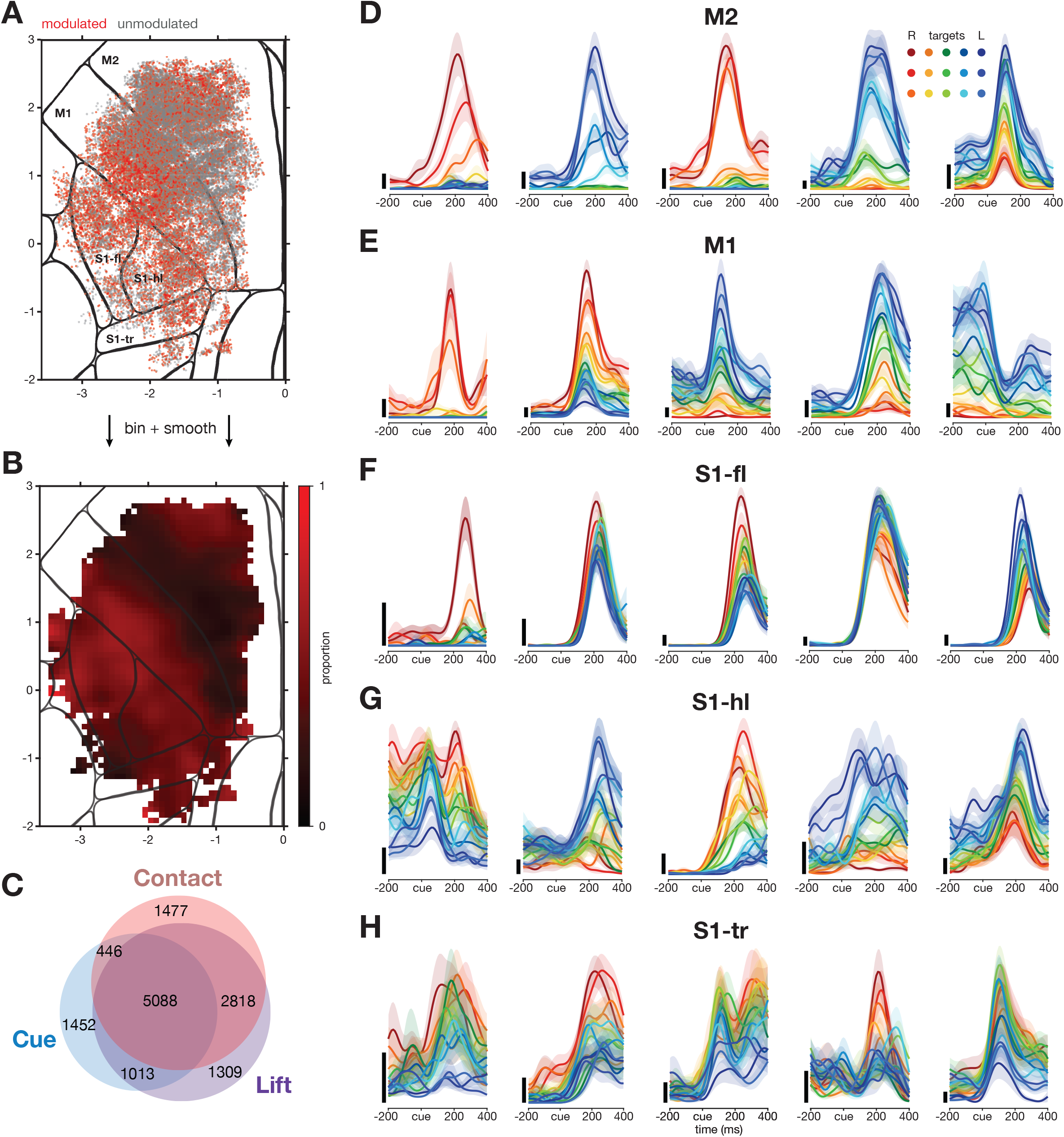
Neurons in mouse sensorimotor cortex exhibited heterogeneous tuning profiles. (A) Extracted neuron locations superimposed on the Allen CCF. Red indicates a neuron that was modulated for any of the three locking events according to our statistical test (Methods), gray indicates a neuron that was not. (B) Binned and Gaussian-smoothed (s.d. 150 µm) map derived from A. (C) Venn diagram depicting the number of modulated cells for each event alignment. (D-H) Peri-event time histograms (PETHs) of five example neurons in each area. Successful trials to each target were averaged and smoothed. Colors as in Figure 2B. Shaded regions, SEMs; scale bars, 10 events/second. Neurons were chosen to be clear and representative examples of response profiles observed in each area.

Modulation data are shown here computed with neural activity aligned to the Go cue. When aligning data to the paw lift (“Lift”) or spout contact (“Contact”), a mostly-overlapping set of cells were identified as modulated (Figure 3C, Figure 3-figure supplement 1). Although more cells were identified as modulated when using the Lift or Contact event alignments, all results in this work were similar across all three possible alignments. Results are shown through the rest of this work aligned to Cue because that alignment best preserved some unique activity structure in M2 (noted below) while preserving all other findings.

We next visualized the tuning profiles of individual neurons through peri-event time histograms (PETHs). Five representative example neurons for each of the five areas of interest are shown (Figure 3D-H). One striking feature of the neural responses was that neurons were often strongly tuned, with large differences in the activity associated with different targets, despite the fact that the targets were relatively close together in physical space. Several systematic differences between the neurons in each area were also prominent, both in these examples and in our qualitative assessment of the recorded population overall. Neurons in both M2 (Figure 3D) and M1 (Figure 3E) were often strongly tuned, with firing rates ranging from no measurable activity for some targets to strong and reliable responses to other targets. M1 neurons typically exhibited briefer responses than M2 neurons, while M2 neurons were often more sharply tuned – that is, strongly active for a smaller number of targets. S1-fl (Figure 3F) typically exhibited the briefest responses, with a consistent peak of activity for most or all targets. S1-hl (Figure 3G) and S1-tr (Figure 3H) neurons had a large variety of response profiles, frequently showing sustained and temporally complex responses and having activity peaks at substantially different times for reaches to different targets. While examples of almost any response profile could be found in each of the recorded areas, these observations reflect systematic differences in the distributions of responses across these areas, quantified below.

### Onset of neural activity varied with somatotopy and subregion

To quantify the overall temporal profile of single neurons, we measured when activity for each neuron ramped up. To compute these onset times, we first averaged the activity for each modulated neuron across all trials, creating a single grand-mean trace per neuron, then found the time to half-max. We next asked whether onset times differed by anatomical area (Figure 4A). These distributions over neurons revealed clear differences in the overall profile of activation: early onsets were more prevalent in S1 trunk and hindlimb regions, perhaps due to activity related to the animal stabilizing itself even if the neurons became more active later; then M2, and finally S1-fl and M1. Nevertheless, each area contained neurons activated at any given time in the trial.

**Figure 4.**
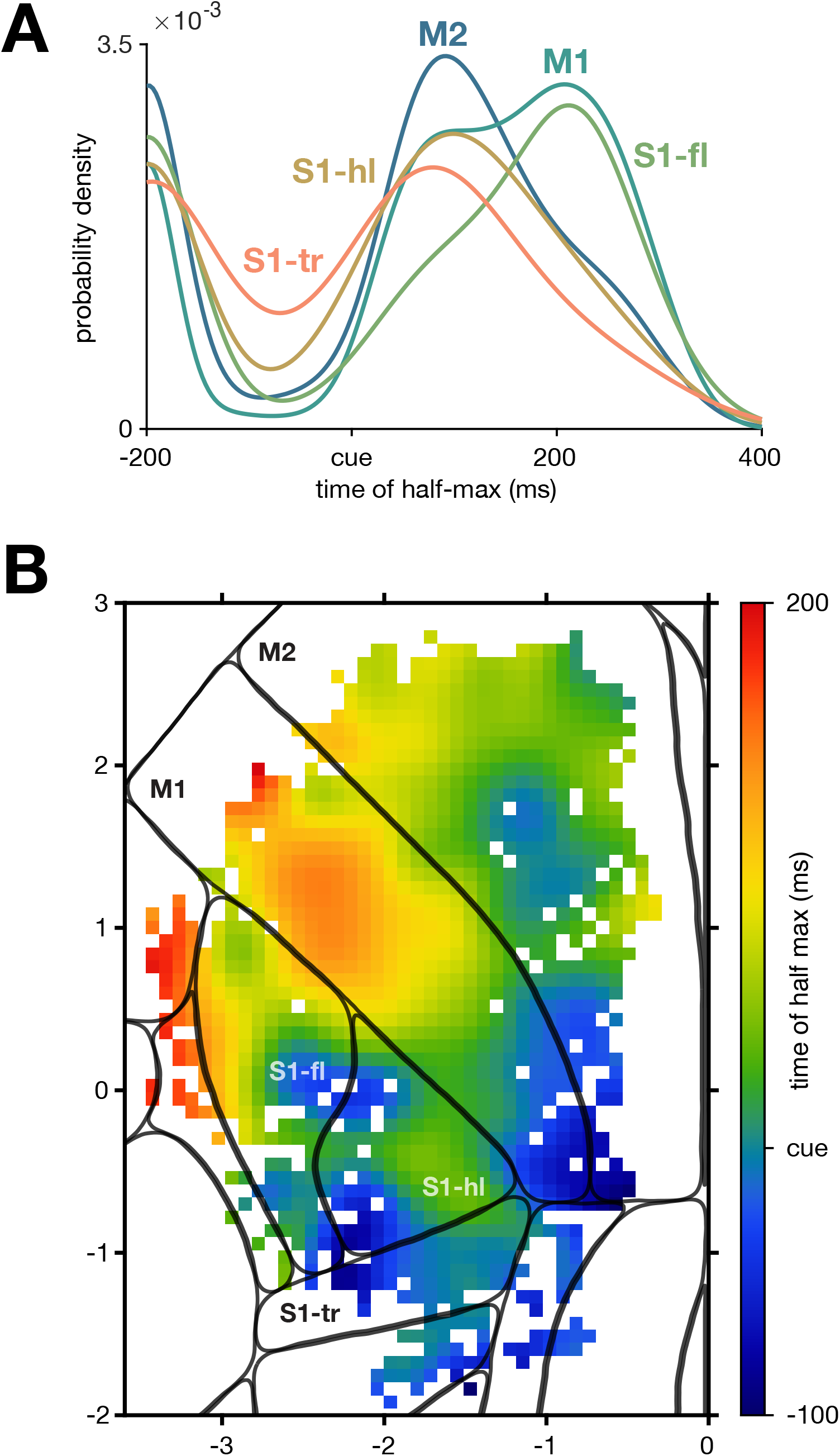
Onset of neural activity across sensorimotor cortex followed both area borders and a somatotopic organization. (A) Smoothed histogram of time to half-max activation for neurons by anatomical area. (B) Time of half-max activation map. The metric was computed on individual neurons, neurons were binned into pixels, then the map was Gaussian smoothed (s.d. 150 µm). Colorbar ticks indicate range of values in binned and smoothed map.

Next, we leveraged our dense spatial sampling to build a map of the time of activation across cortex. The temporal onset metric was computed for each neuron, neurons were binned into pixels, then this map was lightly smoothed (Methods). The resulting temporal onset map revealed a number of sub-area features (Figure 4B). The earliest activations, occurring during the delay (after the target had moved in but well before the Go cue), were in S1 at the juncture of the trunk, hindlimb, and forelimb regions; and in the portions of M1 and M2 that correspond somatotopically to the hindlimb. As suggested by the area-level result above, these zones presumably reflect the animal preparing its body to support the reach. Shortly after, there was activation in a small zone of S1 that corresponded closely with the tactile subregion of the forelimb region, and in the vibrissal subregion of M2 (see Figure 1A; Esmaeili et al. (2021); Mayrhofer et al. (2019)). This latter activation might relate to the animal whisking, possibly to find the target, or because the incoming spout simply triggers a whisking response, or because the animal is engaged (Steinmetz et al. 2019) and anticipating movement. Next, activation spread through much of M2, part of the forelimb portion of M1, and into other parts of S1; and then finally to more lateral parts of M1 and M2 that relate to the face and oromanual feeding (Figure 1A; (An et al. 2025)).

### Activity profiles of single neurons varied systematically across sensorimotor cortex

To quantify the heterogeneity of tuning profiles across single neurons, we developed several summary metrics that captured the most salient variations in observed properties across our inspection of thousands of PETHs. This provides greater interpretability than using unsupervised methods such as Principal Component Analysis (PCA), and is consistent with the practice of feature engineering to extract known-meaningful structure from data where more generic methods fail to capture this structure (e.g., in spikesorting; Caro-Martin et al. (2018), Lee et al. (2021), Lewicki (1994)). Brief intuition is given for each metric below, with complete explanations of each in the Methods.

#### Response duration

To capture whether a neuron was active only briefly or for a more extended time, we measured the width of the autocorrelation for each neuron. Autocorrelations were averaged across all targets for which the neuron was modulated. Larger values indicate that a neuron remained active for longer. This metric was highest in the posterior regions S1-tr, S1-hl, and the portions of M1 and M2 corresponding to the hindlimb (Figure 5A).

**Figure 5.**
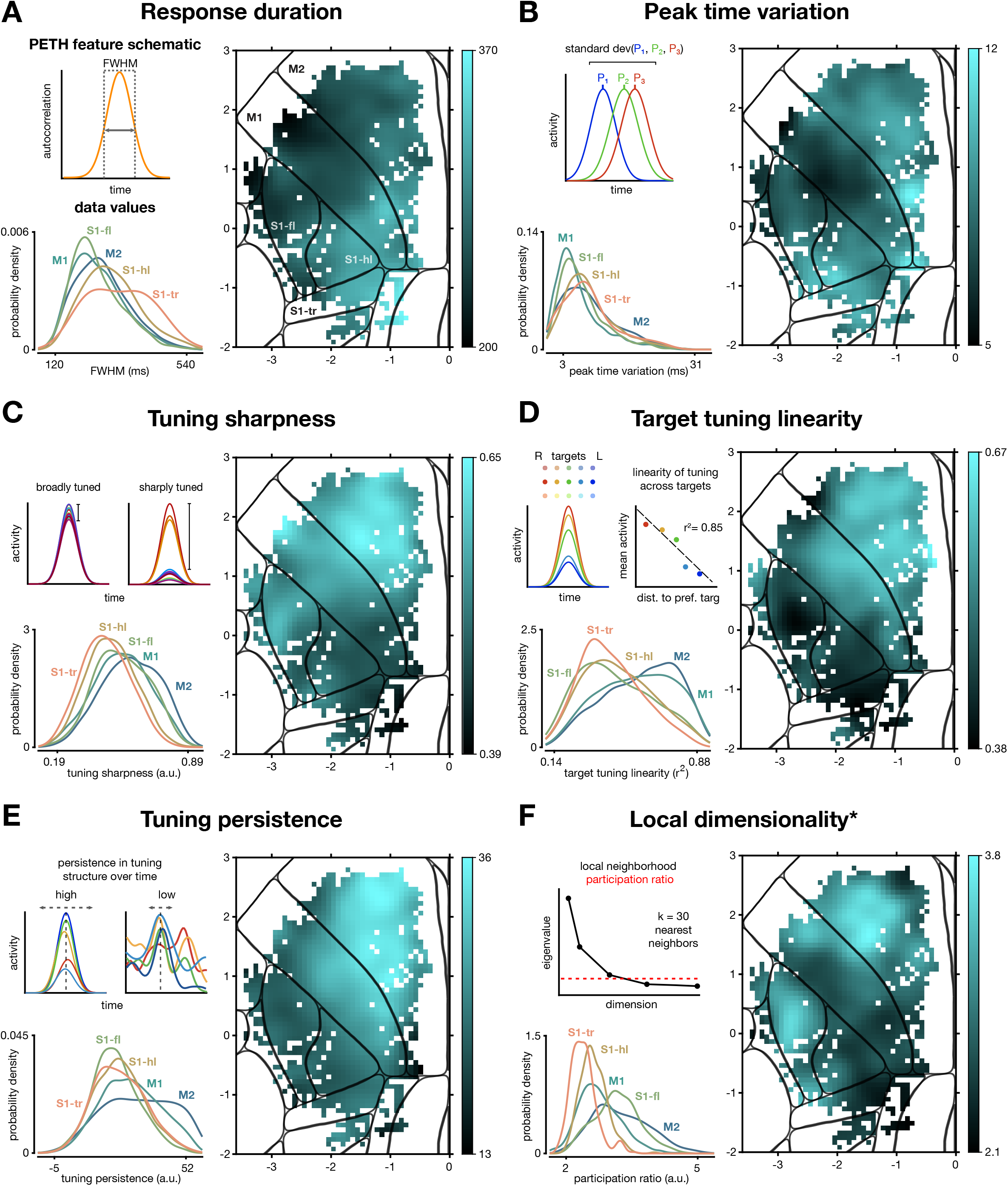
PETH features were organized into distinct spatial patterns in sensorimotor cortex. (A) *Top left*, schematic for the response duration metric, which measures the autocorrelation width of the trial-averaged trace for each neuron and target. *Bottom left*, histogram of response duration values for all modulated neurons grouped based on anatomical region. *Right*, metric map of the response duration values where bright regions correspond to higher response duration values and darker regions to lower. (B-F) Analogous to A for other metrics. (B) Peak-time variation, computed as the standard deviation of peak times across targets. (C) Tuning sharpness, computed as one minus the ratio of the average peak response (excluding the strongest response condition) normalized by the strongest peak response. (D) Target tuning linearity, the coefficient of determination of the peak response with ordinal distance of the target from the target that evoked the strongest response. (E) Tuning persistence, corresponding to how consistent the tuning was across targets at all lags from the peak (Methods). Only statistically-modulated targets were included for each neuron. (F) “Local dimensionality”, the participation ratio of the 30 nearest neighbors to each neuron. Star indicates that this metric was not used for analyses below, because it is not independent for each neuron.

#### Peak time variation

To quantify whether a neuron’s firing peaked at the same time for every target or varied by target, we found the peak firing rate of the response to each target, then computed the standard deviation of these peak times across targets. This value is therefore higher if the peak time varied and nearly zero if the timing was consistent. Notably, this measure correlated substantially with overall signal-to-noise ratio of a neuron’s PETH (Spearman’s ρ=-0.53; Methods), and thus partly measures trial-to-trial variability, not just true peak timing variability. This metric was quite low in M1, indicating highly consistent timing of the activity peak (and reliable responses), and was highest in the posteromedial part of M2 (presumably corresponding to the hindlimb representation) and the posterior tip of S1-hl (Figure 5B).

#### Tuning sharpness

To ask whether a neuron was active for only a few targets or many, we computed a metric we refer to as PETH tuning sharpness (Figure 5C). Neurons attained values near zero if they were similarly active for most targets (i.e., active but broadly tuned), while neurons attained values near unity if they were strongly active for few targets (i.e., sharply tuned). This metric showed that neurons in forelimb-related subregions had sharper tuning than neurons in hindlimb-related regions, with M2 neurons being particularly sharply tuned.

#### Target tuning linearity

To quantify how linearly a neuron’s activity related to target location in physical space, we correlated the 15D vector of mean activity of the neuron for each target with the 15D vector of the targets’ ordinal distances from the neuron’s preferred target (Methods). The resultant metric map shows that the motor areas were much more linearly tuned with respect to target location than were the somatosensory areas, with M2 even more linear than M1 (Figure 5D).

#### Tuning persistence

To capture whether a neuron’s tuning profile was similar across time points or varied at different time points, we computed a tuning persistence metric. Neurons with high persistence maintained the same tuning across longer stretches of time than neurons with low values. Tuning persistence was highest in the anterior of M2, was moderate in M1, and was low in the sensory regions (Figure 5E).

#### Local dimensionality

To ask how heterogeneous small neighborhoods of neurons were, we computed a “local dimensionality” metric: for each neuron, we computed the participation ratio (an imperfect but simple eigenspectrum-based measure of dimensionality) of its 30 nearest neighbors. Higher local dimensionality reflects locations where nearby neurons exhibited more heterogeneity in their PETHs with respect to one another in their dominant patterns (but not necessarily the full diversity of signals present in the population if a few large signals are shared). This measure of local dimensionality was highest in a patchy portion of anterior M2, and in anterior S1-fl (possibly corresponding to the tactile region, see Figure 1A from (Alonso et al. (2023)) (Figure 5F). Note that because this metric involved neighborhoods instead of single neurons, it was not included in subsequent analyses.

We also compared our metric maps with maps generated from the top 20 PCs of the PETHs (Methods), rotated using VARIMAX to identify a sparser basis (Musall et al. 2019). These maps also exhibited some spatial structure, but as expected, when compared with our bespoke PETH features created using knowledge of the problem domain, the spatial structure was weaker (Figure 5-figure supplement 1).

### PETH features changed more strongly at anatomical and somatotopic boundaries

Each of our PETH feature maps varied across the sensorimotor areas, and appeared to change more abruptly in some locations. We wished to ask whether there were in fact relatively discrete transitions in the PETH feature values across cortical space, or whether these values changed only smoothly. To do so, for each PETH feature we computed the gradient: the vector-valued derivative of the PETH feature with respect to two-dimensional cortical space. We then took the magnitude of this gradient computed at every point in the map (Methods; Figure 6A). Intuitively, this simply asks how quickly the PETH feature is changing at each point on the map. Each map revealed a ridge at the edge of the high-value region (Figure 6A-E), suggesting that the PETH feature maps had edges, albeit soft ones, as opposed to smoothly falling off in value. In the target tuning linearity’s gradient map (Figure 6A), this boundary was strongest along the border between M1 and S1-fl, cut through the anterior horn of S1-hl (consistent with projection neuron patterns; Ueno et al. (2018) and Yang et al. (2023)), and separated what is presumably the forelimb representation from both the hindlimb representation posteromedial and the orofacial representation anterolateral. Other PETH features strongly separated M1 and M2 and somatotopic zones (Figure 6B,D,E). Each of the candidate functional boundaries was present in at least two of the gradient maps. We therefore pooled the identified boundaries by computing an overall gradient magnitude (the quadratic mean of the individual PETH feature maps; Figure 6F). This map makes clear that the PETH features of single cells changed most sharply along both area and somatotopic boundaries. Notably, these boundaries did not correspond with ‘edges’ in the map of where more or fewer cells were modulated (Figure 3B).

**Figure 6.**
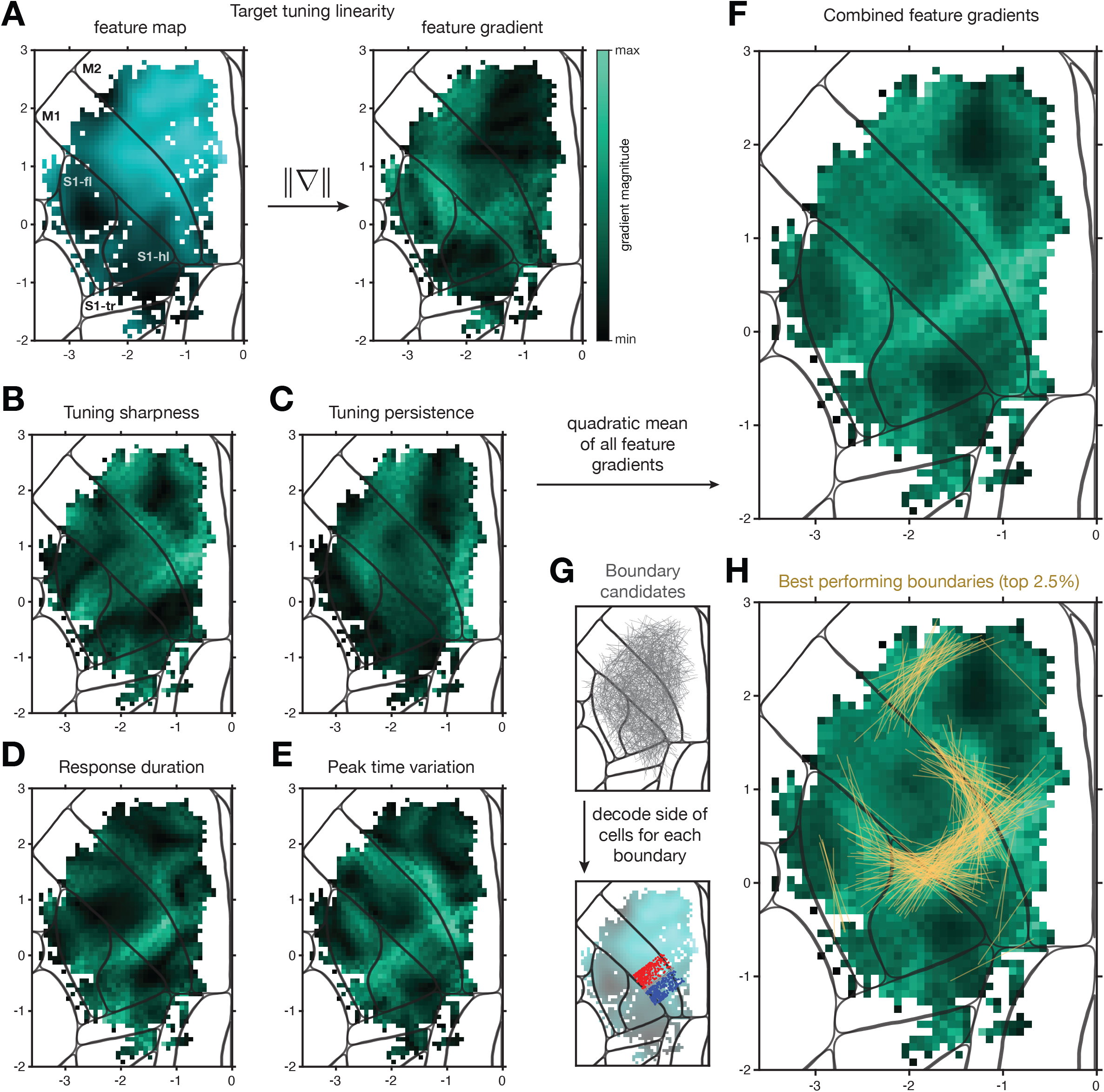
Derivatives of PETH feature maps produced response-property boundaries aligned with anatomy and somatotopy. (A) Side-by-side comparison of a PETH feature map and its gradient magnitude map. *Right*, bright green values represent large gradient magnitudes and dark values represent small gradients. (B-E) Same as A *right*, for the other PETH features. (F) Quadratic mean over the five PETH features of the gradient magnitude maps. (G) *Top*, 10,000 1-mm random boundaries plotted in gray within the sampled region. *Bottom*, for each random boundary, neurons that were within 500 µm on either side of the boundary were included; neurons within 50 µm of the boundary were excluded. The target tuning linearity feature map is shown underneath for illustration. An SVM was trained to predict which side of the boundary a neuron was on using its PETH feature vector. (H) The top-performing 2.5% of random boundaries (assessed via cross-validation) plotted in yellow on top of the average gradient magnitude map.

This gradient method has the advantage of producing a smooth map with simple operations. However, to validate the result, we also used a decoding approach. For each of 10,000 randomly-placed candidate boundaries, we attempted to classify which side of the boundary nearby neurons were on using their PETH features (Figure 6G; Methods). We expected that classification would be better when the random boundary happened to match places where PETH features were changing more rapidly. The top-performing 2.5% of random boundaries are plotted in Figure 6H. They aligned well with the boundaries revealed by the gradient method.

### Areas exhibited distinct and multimodal activity profiles

From the PETH feature maps and distributions (Figure 5), it was clear that areas exhibited distinct response profiles on average. However, within each area, neurons’ responses were heterogeneous. To better understand the distribution of response profiles within and across areas, we used nonlinear dimensionality reduction (t-distributed Stochastic Neighbor Embedding, t-SNE; Hinton and Roweis (2002)). Specifically, we summarized each neuron’s response with a five-dimensional feature vector comprising the five PETH features described above, then applied t-SNE to these vectors for the full neural population spanning all five brain areas (Figure 7A). Finally, we found where the neurons from each area were embedded in the t-SNE space (Figure 7B). The resulting two-dimensional t-SNE projection led to three findings. First, single neurons from each region were found across most of the t-SNE space (Figure 7B). This makes clear that much of the diversity of neuron activity patterns was observed in each area. Second, when looking at where neurons in each area most commonly appeared in the t-SNE space, most areas had peaks centered at different locations – though with overlapping distributions (Figure 7C; all pairwise comparisons p<0.005, two-dimensional Kolmogorov-Smirnov test). The largest differences were between motor and somatosensory areas, with S1-fl bridging the other S1 subregions and M1. This implies that areas had distinct distributions of their PETH features, but also that there was an interpretable spectrum of typical properties from M2 to M1 to forelimb S1 to hindlimb and trunk S1. Third, the distributions of PETH features within-area were strongly multimodal for several areas. This suggests that the heterogeneity of neurons within each area may form loose response ‘types,’ especially in M2, M1, and S1-fl. This multimodality motivated further analysis targeted at understanding the range of neural response profiles in each area, and whether neurons with specific response profiles were preferentially found in localized regions of each cortical area.

**Figure 7.**
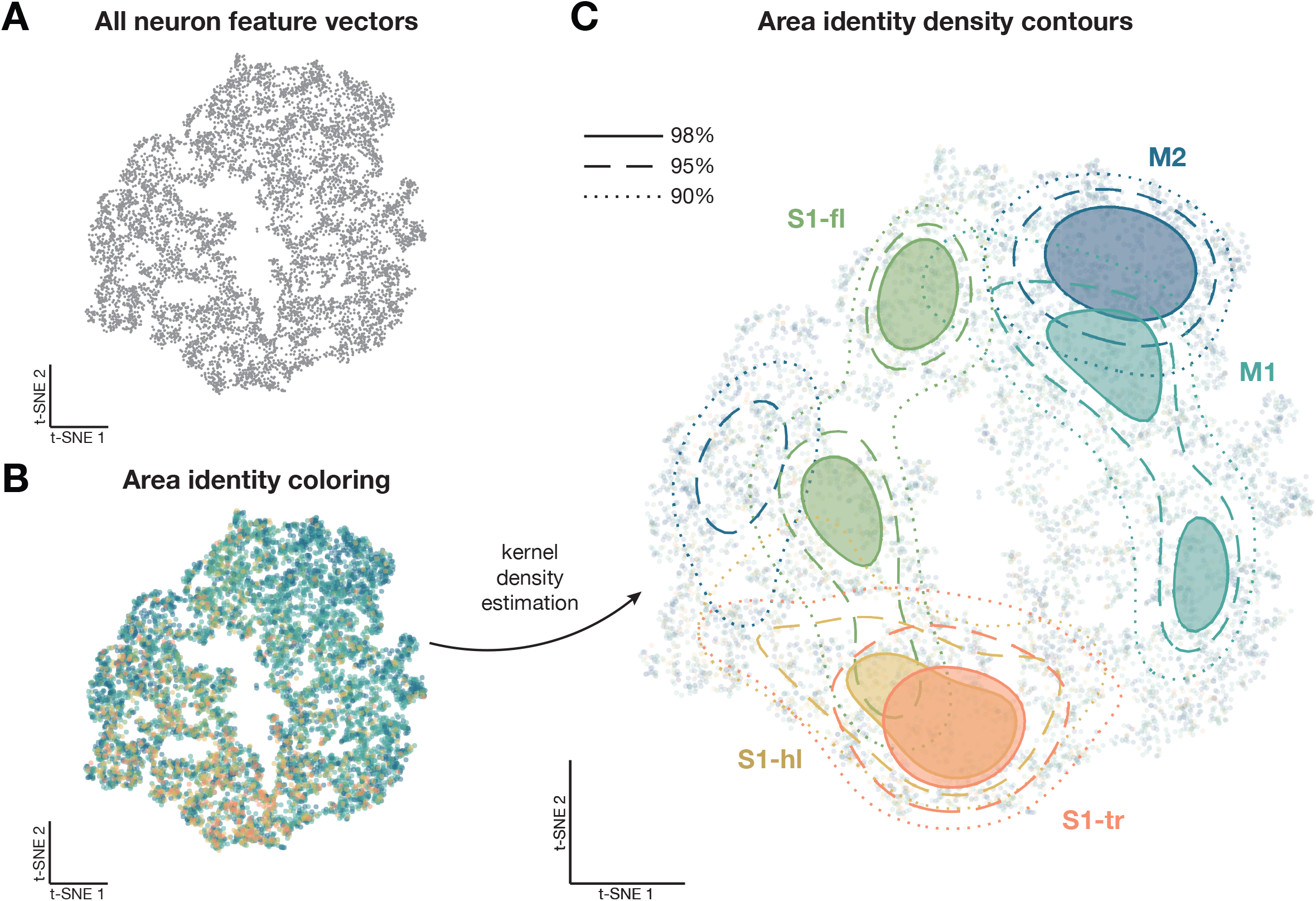
Areas had distinct and multimodal distributions of PETH features. (A) t-SNE of the five-dimensional PETH features for all neurons. Each gray point corresponds to a neuron. (B) Same embedding as in A, with neurons (points) colored by anatomical area. (C) Contour plots of the embedding from B, using two-dimensional Kernel Density Estimation to produce a smooth density estimate. Three contour levels are shown for each of the five anatomical areas of interest.

### Neighboring sensorimotor regions shared PETH features in complex spatial patterns

The multimodality of feature vector distributions within-area opens a range of possibilities for how PETH response features could be organized across the cortex, both within and across areas. To work with these multimodal distributions, we fit a generative model to the PETH feature vectors defined above for each area’s population of neurons. This approach modeled the feature vectors of an area’s neurons as arising from a Gaussian Mixture Model (GMM). Doing so explicitly models different peaks in the distribution as arising from different Gaussian distributions, and retains probabilistic interpretability. In agreement with the t-SNE results, M1, M2, and S1-fl were best modeled by the GMM as having multiple components (Methods): four for M2, three for M1, and two for S1-fl, while S1-hl and S1-tr were best modeled as being unimodal. The Gaussians fitting different modes were well-separated, as expected when fitting a multimodal distribution (Figure 8-figure supplement 1).

We hypothesized that even though the distributions of response profiles were multimodal within-area, different areas might still be well separated. To test this, we computed a data likelihood for each neuron – p(neuron *i*’s PETH features | GMM_area_) – as arising from each area’s distribution (Figure 8A). Intuitively, this produces a measure of how similar each neuron is to the population of neurons in an anatomically-defined area. We then binned and smoothed over neurons by cortical location to produce the maps in Figure 8B-F. This mapping approach is explicitly biased toward finding feature differences between areas, allowing for a direct test of the hypothesis that response profile distributions are area-specific.

**Figure 8.**
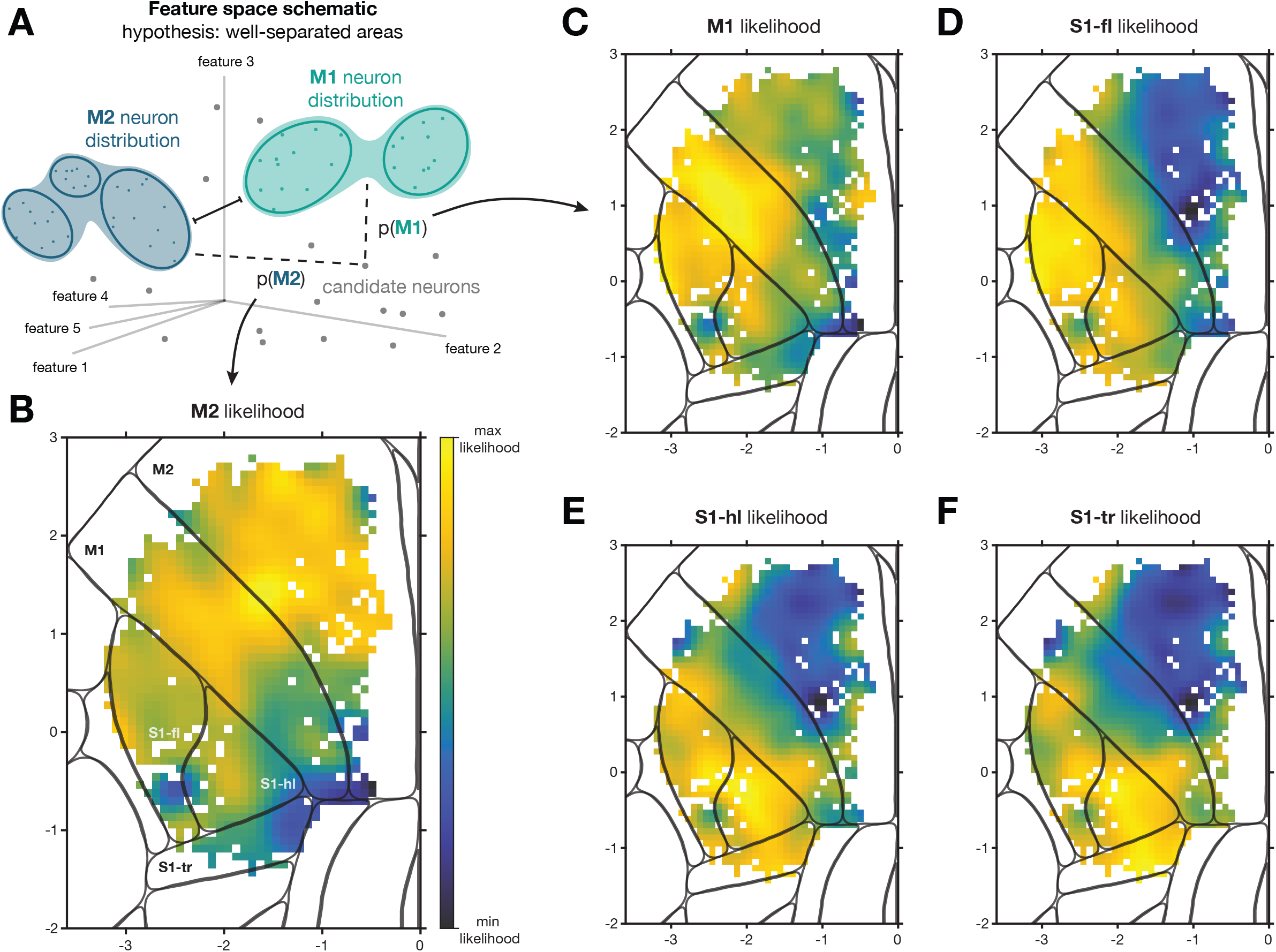
Neural response profiles were shared in patches that spanned anatomically defined regions. (A) Schematic of analysis with hypothesis that neurons from different areas are well-separated in feature space. A Gaussian Mixture Model (GMM) was fit to the distribution of feature values for neurons belonging to a single anatomically-defined area, then a likelihood was computed for all neurons from all areas using the fit. (B) The resulting likelihood map for M2, where bright yellow pixels correspond to high likelihood regions and dark blue pixels to low likelihood regions. Colorbar scale is logarithmic. The ends of the colormap were set to the maximum and minimum likelihood values for each map. (C-F) Same as B, where the seed regions were M1, S1-fl, S1-hl, and S1-tr, respectively.

When computing this likelihood map using neurons in M2 to define the distribution (Figure 8B), the highest likelihood zone was, as expected, in M2. However, we found that a patch of M1 was strongly similar. If we instead defined the distribution using M1 neurons (Figure 8C), again the highest-likelihood zone was within the defining area. However, two additional features were salient. First, the anterior portion of S1-fl was also strongly similar to M1, in line with previous results (Grier et al. 2026). Second, the posteromedial portion of M1 differed strongly from the rest of M1, consistent with a difference between forelimb and hindlimb somatotopic regions (Maurer et al. 2023). Considering other areas, defining the distribution with S1-fl led to high likelihoods in adjacent M1 and S1-hl (Figure 8D), while defining the distribution with S1-hl or S1-tr produced similar spatial layouts and bled over mostly into the other sensory areas (Figure 8E,F). These results make clear that the working hypothesis – of areas with well-separated feature distributions – is incorrect. Even when building models explicitly from neurons in one area, neurons in portions of some areas were similar to neurons in portions of other areas, and two dissimilar areas could both be similar to the same third location in cortex. These neural populations therefore warranted further dissection.

### Subpopulations spatially overlapped and spanned anatomical areas

The likelihood maps above suggested that populations of neurons with similar response profiles were not always confined to the anatomically-defined areas. For example, portions of M1 and S1-fl were strongly similar (Figure 8BC,D). This hinted at a second hypothesis: that a peak in the multimodal distribution from one area might correspond to a peak in the multimodal distribution of a different area. If this were due to a subpopulation that was shared across the areas, we would expect that the M1 GMM would contain a component (a one-area estimate of the subpopulation) with similar characteristics to a component in the S1-fl GMM (Figure 9A). We therefore computed a distance metric (Methods) between each pair of GMM components (Figure 9B). This analysis revealed that many components identified independently in different areas were indeed highly similar, and formed four clusters. We therefore combined components within each of the four clusters (Methods) and took the neurons belonging to each cluster as the subpopulations.

**Figure 9.**
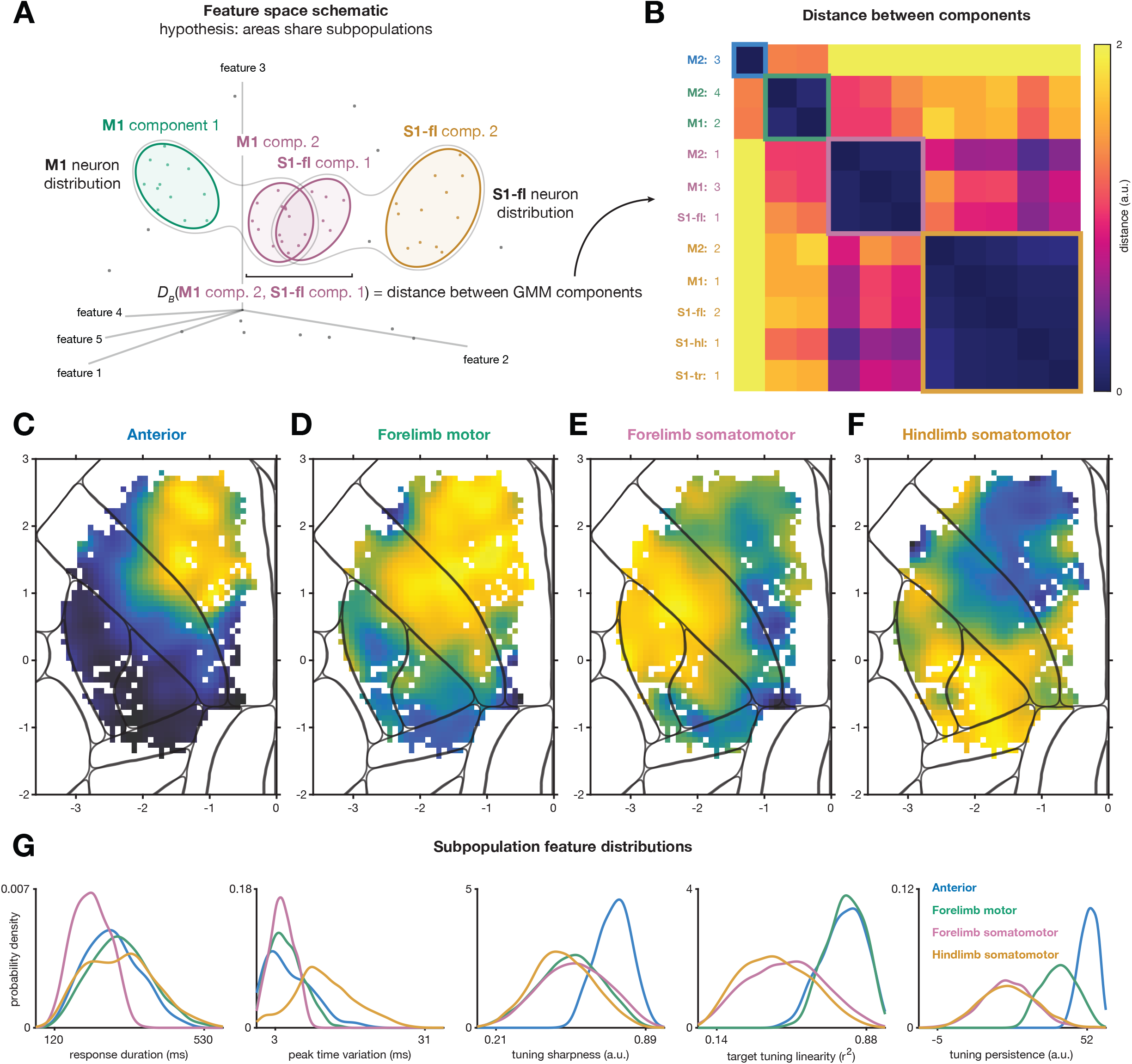
Four subpopulations with distinct response profiles form an overlapping patchwork across sensorimotor cortex. (A) Schematic of GMM modeling with the hypothesis that components from different areas correspond. Only two GMM components in each area are illustrated for simplicity, with numbers corresponding to those in B. (B) Bhattacharyya distance calculated between all pairs of GMM components across all anatomical regions. The matrix is organized based on cluster identity from hierarchical clustering of the pairwise distances. Each cluster (colored box outlines) is referred to as a subpopulation. Colormap capped to better show any within-cluster detail. Cue-locked data, modulated neurons only. (C) Anterior subpopulation likelihood map. Plotting as in the other likelihood maps with colormap as in Figure 8. (D-F) Same as C but for Forelimb motor (D), Forelimb somatomotor (E), and Hindlimb somatomotor (F) subpopulations. (G) Distributions of the five feature values (subpanels) for each of the four subpopulations (traces)

These four subpopulations differed in two major ways. First, by construction, different subpopulations were characterized by distinct PETH features (Figure 9G). Second, their spatial likelihood maps differed strongly as well (Figure 9C-F). One subpopulation, which we refer to as the “Anterior” subpopulation, was almost unique to M2 (Figure 9C). It had uniquely high tuning persistence and tuning sharpness along with highly linear target tuning, arguing for a relatively abstract and external target representation (Galiñanes and Huber 2023). A second subpopulation, the “Forelimb motor” subpopulation, spanned M2 and the forelimb portion of M1 (Figure 9D). These neurons were also strongly linear in their target tuning, but less sharply tuned and with tuning that was somewhat less persistent (i.e., changed more over the course of the movement).

The “Forelimb somatomotor” subpopulation spanned M1 forelimb and orofacial zones and the anterior portion of S1-fl (Figure 9E). This subpopulation was more broadly tuned, was less linearly tuned to target location, changed tuning over the course of the movement (low tuning persistence), and responded very briefly (low response duration). Nevertheless, each neuron in this subpopulation still responded at its own characteristic time in the movement (low peak time variation). Finally, a “Hindlimb somatomotor” subpopulation (Figure 9F) was centered on S1-hl and spilled into S1-tr, the posterior portion of S1-fl, and the posterior tips of M1 and M2. This subpopulation was similar to the Forelimb somatomotor subpopulation, but each neuron no longer fired at a specific time point in the movement; instead, neurons fired at different times for different targets (high peak time variability). As a qualitative depiction of the response profiles identified with each subpopulation, we plotted the two highest-likelihood cells for each area/subpopulation combination (Figure 9-figure supplement 1). These examples reveal stereotypy in the subpopulation responses across areas, but also show variation across areas, especially for the two somatomotor subpopulations.

To better understand whether the subpopulations were well separated in feature space or not, we examined our clustering further using two approaches. First, for each pair of subpopulations, we trained a logistic regression classifier to separate the 5D feature vectors of the neurons in one subpopulation from the feature vectors of the neurons in another subpopulation, then projected the feature vectors onto this axis (Figure 9-figure supplement 2A). The resulting distributions were well separated along this axis in all but one of the pairwise comparisons (Hindlimb somatomotor vs. Forelimb somatomotor subpopulations). Second, we re-examined our t-SNE of the features, contouring density by subpopulation membership instead of area membership (Figure 9-figure supplement 2B). Despite t-SNE being a quite different nonlinear operation than applying a GMM, subpopulations were again distinct. Interestingly, here the strongest separation was between the Hindlimb somatomotor subpopulation and all other subpopulations. To assess an orthogonal property, we examined the distributions of activity onset times by subpopulation (Figure 9-figure supplement 2C). These distributions appear to be more distinct for the subpopulations than for the areas (compare with Figure 4A). Overall, these analyses support the interpretation that the subpopulation distributions are distinct in feature space.

To confirm that this identification of area-spanning subpopulations was not simply due to our choice of PETH features, we validated these findings by repeating some of our analyses using the top 20 PCs of the PETHs instead of the PETH feature vectors. Using PCA as a preprocessing step in this way is more common in the field. Performing t-SNE on the PCs, analogously to Figure 7, again yielded multimodality (Figure 9-figure supplement 3A). Repeating the analysis of Figure 8 on the PCs produced likelihood maps that were substantially patchier and less distinct between areas, but recapitulated that parts of each area were strongly similar to parts of other areas (Figure 9-figure supplement 3C-G). This indicates that although our choice of PETH features enabled a sharper view of the distinctions between subpopulations, these features were not necessary to detect them.

To determine whether finding distinct subpopulations required recordings from a large area, we directly examined a smaller patch of M1 (480 µm diameter) in an overlap zone (Figure 9-figure supplement 3B, *leftmost*). Applying t-SNE to either our PETH features (Figure 9-figure supplement 3B, *left*) or to the top 20 PCs of the PETHs (Figure 9-figure supplement 3B, *right*) again yielded multimodality. This demonstrates that even with a single ordinary-size field of view, distinct subpopulations in overlapping zones could be readily detected in the context of our behavior.

To validate the clustering itself, we fit a GMM to all neurons without consideration of their area membership. This process produced three components with maps that were nearly identical to three of the four subpopulations identified with the area-aware method (Figure 9-figure supplement 4). The only noticeable difference was that the Anterior subpopulation was lost when all areas were fitted together, likely due to having a lower membership than the other subpopulations. Using this area-agnostic model we also demonstrated that these three main subpopulations were reliably detected in subsets of our data. To do so, we evaluated how similarly two GMMs fit to 50/50 area-wise splits of the data clustered the neurons. The median Adjusted Rand Index (Hubert and Arabie 1985), a measure of clustering similarity, was 0.856 across splits, indicating highly similar clusterings from the data subsets. Together, these controls indicate that the clustering was reliable.

Notably, the high-likelihood regions for different subpopulations spatially overlapped. The Anterior and Forelimb motor subpopulations overlapped throughout M2; the Forelimb motor and the Forelimb somatomotor subpopulations overlapped in the forelimb portion of M1; and the Forelimb somatomotor and Hindlimb somatomotor subpopulations overlapped in a band along the S1-fl and S1-hl border. This indicates that although several of the PETH feature boundaries found in Figure 6 aligned with area borders, several of these boundaries corresponded to the edges of subpopulations that spanned other borders. To confirm this interpretation, we computed the gradients of the likelihood maps directly (Figure 9-figure supplement 5). These maps closely matched those computed from the PETH feature gradients directly (Figure 6), as expected.

### Subpopulation members were intermingled and reflected similar response profiles across areas

The overlaps in the subpopulation likelihood maps above imply that members of different subpopulations are spatially intermingled, but it is less clear whether each subpopulation has homogeneous response profiles across space. In particular, the use of likelihoods mixes two properties: the fraction of neurons in a given neighborhood that are members of each subpopulation, and the heterogeneity of response profiles amongst members of that subpopulation. These properties could vary systematically with respect to one another, and the spatial structure shown by the likelihood map does not disentangle them.

We therefore assessed the distribution of subpopulation memberships across cortex independently from response profile variability within subpopulation. To do so, we first classified each neuron into its maximum-likelihood subpopulation, then mapped the subpopulation identity of neurons (Figure 10A). As expected, members of the different subpopulations were locally salt-and-pepper intermingled, including in overlap zones. Also consistent with the likelihood maps, the fraction of each subpopulation in each area differed (Figure 10B). Critically, we found that the prevalence of subpopulation membership across cortex matched the likelihood maps (Figure 10D), implying that the spatial structure in the likelihood maps reflects the spatial variation in prevalence of each subpopulation and not some other structure in the heterogeneity of response profiles.

**Figure 10.**
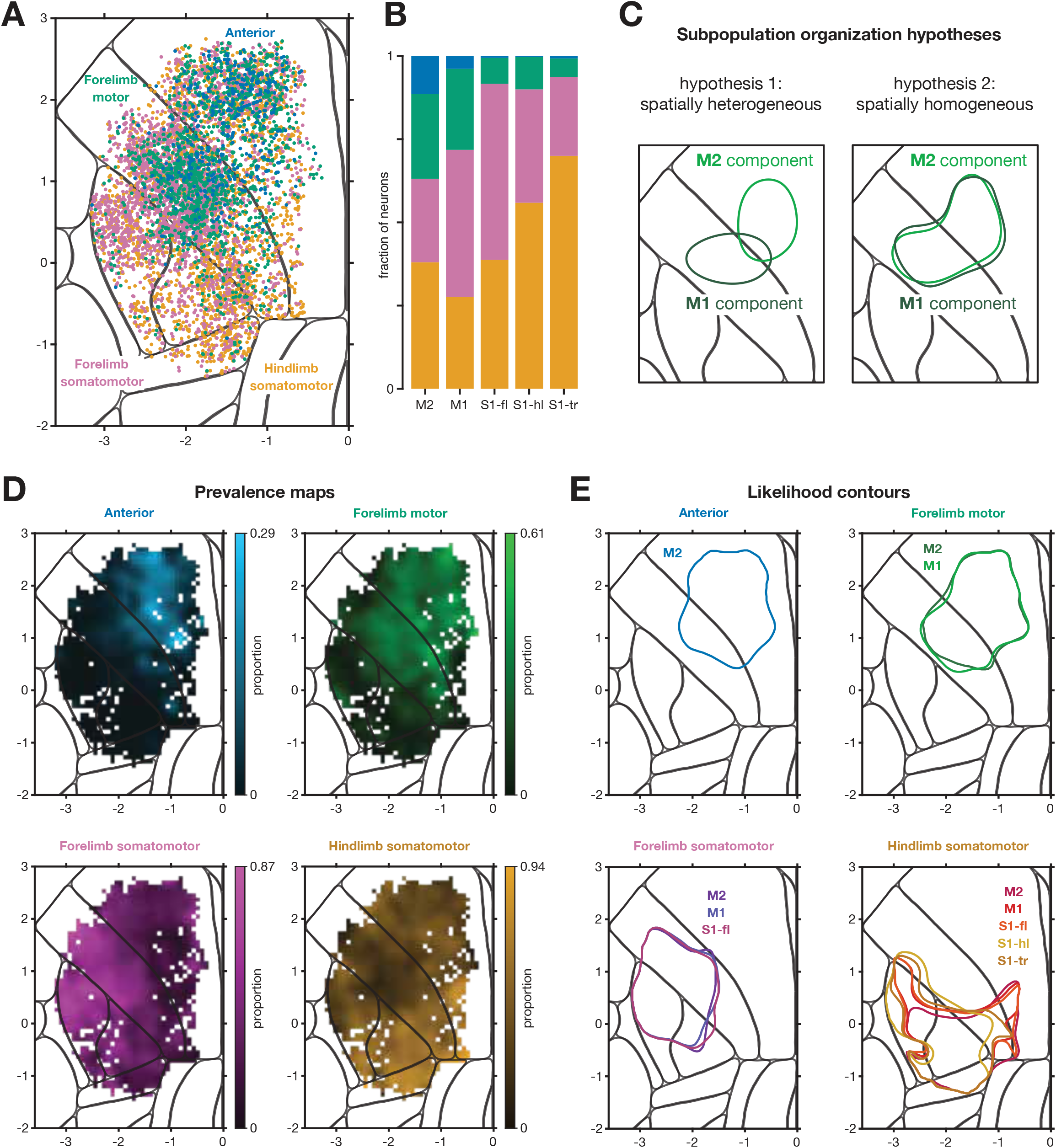
Subpopulations overlapped with their member neurons spatially intermingled. (A) Neurons were classified into their most likely subpopulation, color coded and plotted on the CCF. (B) Prevalence of subpopulation by area. (C) Schematic for distinguishing two hypotheses, if subpopulations were identified separately using data from each of two areas and the identified high-density zones of neurons were plotted as contours of the likelihood maps. *Left*, expected result if subpopulations were homogeneous in their properties over cortical location; *right*, if subpopulations varied smoothly with cortical location. (D) Maps of prevalence of each subpopulation. (E) Likelihood map contours at 60% level, for each subpopulation identified separately using data from each area containing sufficient subpopulation members.

Finally, we looked for heterogeneity in the PETH features within the footprint spanned by each subpopulation. If the PETH features within a subpopulation varied substantially across areas, a model built on neurons from one area (e.g., M2) would assign lower likelihoods to neurons in other areas (e.g., M1), and vice versa (Figure 10C). To determine if this was the case, we produced likelihood maps for each GMM component of the models fit to each area separately, thresholded these maps to produce a contour, and compared their shapes. Subpopulations had nearly identical spatial footprints no matter which area was used to identify them (Figure 10E). This was true even in cases where only a small fraction of neurons in an area were members of that subpopulation: for example, the Forelimb somatomotor and Hindlimb somatomotor subpopulations could be identified using M2 data, even though there was no high-likelihood zone for these subpopulations within M2 (see Figure 9E-F). Note that this consistency mirrors the low Bhattcharyya distances between corresponding GMM components in Figure 9B, and further validates the use of clustering to combine components from different areas. Together, these results argue that the overlap zones for different subpopulations contained locally intermingled members, and that the PETH features within each subpopulation were similar across the territory covered by that subpopulation.

## Discussion

By densely sampling layer 2/3 of mouse sensorimotor cortex during a 15-target reach-to-grasp behavior, we mapped how single cell response characteristics were distributed across five areas. Neurons responded heterogeneously within all imaged areas, and within M2 and M1 in particular, but the average response properties of areas were distinct. In parallel, we identified four subpopulations of neurons with unique activity profiles, which spanned multiple areas and were spatially intermingled in the zones where different subpopulations overlapped. One subpopulation was unique to M2, one spanned M2 and the forelimb portion of M1, one was centered in S1-fl but extended into S1-hl and the forelimb and orofacial portions of M1, and one spanned the likely hindlimb representations of S1, M1, and M2. Members of these subpopulations did not vary substantially with their spatial location, suggesting that the average differences observed between areas were largely, though probably not exclusively, a function of their containing different proportions of the subpopulations. These activity-based maps thus reveal new structure in sensorimotor cortex and complement a variety of existing mapping methods.

The investigation of coding here was somewhat different than the traditional approach. We did not focus on traditional questions of “what” variables are encoded, such as external target location vs. kinematics or reach direction vs. reach duration. Instead, we focused on “how” task-related activity was structured, such as response duration, timing of peaks across targets and tuning linearity. This feature-engineering approach produced a complementary picture to the more common unsupervised approach to feature extraction, using PCA on the PETHs (as used previously; (Esmaeili et al. 2021)). This makes clear that there are potentially important differences not just in the content of the activity (e.g., encoding target vs. movement commands (Grier et al. 2026)), but also its format (e.g., linear encoding vs. nonlinear, persistent vs. brief responses). In particular, we found that anterior areas tended to have responses that were sharply tuned for target and preserved throughout the movement, while posterior areas tended to be broadly tuned for target and exhibited greater temporal heterogeneity.

What, then, might be the functional roles of these subpopulations? While future work relating neural responses to behavior will be essential to answering this question, the present analyses suggest some possibilities. M2 contained neurons that were sharply and linearly tuned for target location, as well as having earlier onsets, longer responses, and consistent tuning over the movement. This might suggest a more abstract (Bernardi et al. 2020) or external encoding (Galiñanes and Huber 2023). S1-hl and S1-tr also had early responses, with activity spanning the movement but differing in complex ways across targets. This may suggest activity related to bracing or stabilizing the animal (Disse et al. 2023; Nandakumar et al. 2021). The responses of S1-fl were briefest and typically weakly tuned for target, but peaked at a tightly-timed moment in the movement for most or all targets, with the peak timing being neuron-specific. These responses may correspond to some specific sensory signal like afferent feedback during paw lift (Grier et al. 2026; Mathis et al. 2017), but understanding what that signal would be will require future study. The shape of the Forelimb motor subpopulation is similar to the shape of a recently proposed organization based on corticofugal projection patterns, the Medial Anterior Cortex (MAC) (Yang et al. 2023). This would be consistent with MAC’s proposed role in reaching. Finally, M1 was at the center of the overlap between subpopulations. This is reminiscent of its suggested role in taking top-down commands from M2 and elsewhere and bottom-up sensory input from S1 to actively control the limb (Galiñanes and Huber 2023; Heindorf et al. 2018; Terada et al. 2022). In each of these cases, determining the relationships of the observed activity patterns to function will require specific attempts to link the activity to kinematics, target location, sensory feedback, and more; these relationships will be addressed in future work.

The area-spanning nature of the subpopulations could arise from a number of possible organizations of neural circuits. One hypothesis is that each subpopulation represents a single recurrent circuit that is distributed across areas. In this hypothesis multiple circuits exist in parallel within each area, each strongly connected within the subpopulation and less strongly connected with the other subpopulations. An alternative hypothesis is that a single area contains the circuit that generates the signals characteristic of a single subpopulation, then this activity is broadcast to other areas through the strong connectivity of sensorimotor cortex (Muñoz-Castañeda et al. 2021; Zingg et al. 2014). This broadcast would then cause that characteristic activity pattern to appear in the recipient areas. Recording neurons that project from one area to another, and recording from other layers corresponding to later stages of processing and outputs, will be critical to distinguishing these hypotheses and more fully understanding these circuits.

The maps produced here, using response properties of neurons during task performance, aligned with anatomical and stimulation-based maps in some respects but differed in others. We identified response-property boundaries that aligned closely with the anatomical borders between areas M2, M1, and S1. We also identified response-property boundaries consistent with somatotopy, separating forelimb from hindlimb and forelimb from orofacial representations. Interestingly, these somatotopic boundaries were generally stronger than those corresponding to “true” area borders, despite the known coarseness of somatotopic maps in primate motor areas (Mitz and Wise 1987; Neafsey et al. 1983; Qi et al. 2010; Van Acker et al. 2013); whether the maps are similarly coarse in mouse remains to be tested. Some of our maps also distinguished the tactile portion of S1-fl from the remainder of the area, and grouped the proprioception-receiving portions of S1-fl and M1 (Alonso et al. 2023). The spatial distribution of modulated cells in Figure 3 suggests a distinction between the caudal forelimb area (CFA, involving M1 and S1-fl) and the rostral forelimb area (RFA) in M2, while the feature gradient boundaries suggest a distinction between M1 and M2 more generally. The absence of a clearly delineated RFA was surprising, given its distinct projection patterns (Carmona et al. 2024; Hira et al. 2013b; Wang et al. 2018) and functional differences from CFA (Kristl et al. 2025; Morandell and Huber 2017; Saiki-Ishikawa et al. 2025), but our results might suggest that the activity in layer 2/3 of RFA does not differ markedly from other nearby subregions of M2. The difference in spatial footprints of the Forelimb motor and Hindlimb somatomotor subpopulations also loosely corresponded to the spatial distribution of projections to the lateral rostral medulla (denoted the lateral anterior cortex or LAC in (Yang et al. 2023)), but assessing exact correspondence will require methods not employed here. Finally, our approach did not clearly identify the anterolateral motor area (ALM; (Chen et al. 2017; Inagaki et al. 2018; Li et al. 2015)) within M2, though this may be a consequence of the time period analyzed; a time window targeted to the period of withdrawal and water consumption might more clearly delineate this region. These various anatomical distinctions might appear in activity patterns during other tasks that require different coordination between subareas, with analysis of other time periods, with use of additional response features, or with recordings from deeper layers. Identifying such functional differences is left to future work.

Crafting additional PETH features, or using end-to-end neural network approaches to discover other features, might enable the discovery of additional structure (Minderer et al. 2019; Wang et al. 2023b). For example, our PETH features were chosen to be invariant to the onset time of activity, but these onset times were markedly later in lateral M1 than in adjacent M2 or S1-fl. Including onset times, using a wider window of time that includes more of the reward/licking period, aligning data to other behavioral events, or adding other PETH features would presumably result in finer subdivisions of sensorimotor cortex. This suggests that although our current methods find a large amount of structure in sensorimotor cortex, more structure in the responses surely exists. Relatedly, using other tasks might reveal additional functional differences, or minimize ones we identified. Similar explorations of data from other behaviors is likely to shed additional light on the organization of responses in these areas.

Our findings of larger-scale structure, in the form of area-spanning subpopulations, may intersect with three prior findings. First, several widefield calcium imaging studies have also found that S1-fl activity groups with activity in portions of M1 and S1-hl (Costa et al. 2025; Musall et al. 2019; Saxena et al. 2020; Zatka-Haas et al. 2021). Relatedly, orofacial representations share similar relationships with behavior across M2, M1, and S1 (An et al. 2025; Musall et al. 2019); and both these spatial patterns of coactivation reflect anatomical connectivity (Zingg et al. 2014). These previous findings of larger-scale grouping are consistent with our results. Second, in several cortical areas, populations with different projection targets have known functional differences (Chen et al. 2013; Currie et al. 2022; Economo et al. 2018; Hwang et al. 2019; Kim et al. 2018; Li et al. 2015; Park et al. 2022). However, these specific populations reside mainly in layer 5, while imaging here was performed more superficially. Third, the existence of subpopulations of neurons in sensorimotor cortices have been found independently by using genetic identification of excitatory neuron subtypes, and these subpopulations exhibit distinct activity patterns (Hira et al. 2013a; Li et al. 2024; Mohan et al. 2023; Musall et al. 2023). Whether those subpopulations correspond to the ones found here cannot yet be reconciled. However, many of the genetically-identified neuron subtypes in those studies are present only in deeper cortical layers, while the imaging performed here was exclusively superficial. This suggests that the subpopulations we have identified are unlikely to correspond closely with the particular subtypes found thus far via genetic tools.

The distributions of PETH properties we observed were multimodal, but these distributions were continuous; we did not observe tight clusters in feature space separated by gaps. In interpreting the observed multimodal distribution, we note that a uniform distribution can be transformed into a multimodal one, or vice versa, by a continuous nonlinear function. This is a fundamental problem whenever searching for structure in the distribution of neural activity: even examining the raw spike rate of a neuron involves the spike threshold nonlinearity, and what property counts as the ‘fundamental, linear’ one is almost a philosophical question. Nonetheless, the potential for nonlinear measurement makes it challenging to determine whether there are true “categories” of neurons (Kaufman et al. 2022; Posani et al. 2025). If the interpretation of separate modes as subpopulations were incorrect under some relevant nonlinear transformation, our results would nevertheless imply that M2 has the highest diversity of responses, with shrinking variety toward the posterior as well as changes in the average response profile. If this alternate interpretation were the strongest conclusion possible, it would still be essential to understanding the layout of function across sensorimotor cortex. Confirming that these modes are more deeply meaningful will require future external validation.

Several limitations of the present work should be noted. First, the data here include only superficial layer 2/3. Activity patterns in other layers, or even in the deeper portion of layer 2/3, may differ (Currie et al. 2022; Heindorf et al. 2018; Masamizu et al. 2014). Second, while we took great care in aligning our FOVs to the Allen CCF, we did not analyze the anatomy of fixed tissue for each mouse. Limitations in our alignment, differences between the anatomy of the strain of mice used here from the strain used for building the atlas, and possible mouse-to-mouse variation in the borders of areas all introduce jitter. The sharpness of the boundaries we identified therefore serves as a lower bound. Third, although we achieved coverage of a large swath of sensorimotor cortex, we sampled motor areas more densely than sensory and not every area was densely sampled in every mouse. We therefore may have underestimated heterogeneity in S1, which was sampled less. Fourth, although we obtained high-speed video of the behavior, this work does not consider the kinematics. The relationship of these neural data to the detailed kinematics will be treated at length in future work. Fifth, this work considered only trial-averaged responses, not single trials. This choice enabled us to compare much larger areas of cortex by pooling across sessions, but precluded many types of analysis including of noise correlations (Kiani et al. 2015; Kohn et al. 2016; Zohary et al. 1994). Future work could leverage simultaneous recording of multiple areas to investigate single-trial co-fluctuations in neural activity within subpopulations and across areas.

In summary, these findings provide a response-based atlas of sensorimotor cortex and reveal overlapping subpopulations with intermingled members. The nature of the circuits that underlie these subpopulations, and the content of communication between areas more broadly, remain questions of fundamental importance for understanding how movement is generated and feedback incorporated.

## Methods

### Subjects and surgical procedures

All procedures were approved by the University of Chicago Institutional Animal Care and Use Committee. For two-photon imaging, six male mice aged 8-12 weeks were used. Mice were crosses of Slc17a7-IRES2-Cre-D (JAX strain 037512; Harris et al. (2014)) and TIGRE2-jGCaMP8s-IRES-tTA2 (JAX strain 037719; (Sweeney et al. 2025)) which express GCaMP8s in most glutamatergic neurons (Wang et al. 2023a). Each animal underwent a single surgery, then were individually housed on a reverse 12-h light/dark cycle, with an ambient temperature of 22 °C and a humidity of 58%. Experiments were conducted in the afternoon, during the animal’s dark cycle.

Anesthesia and basic surgical preparation were as described in (Grier et al. 2026). A circular craniotomy was outlined with a 5-mm biopsy punch and completed with a dental drill, centered over the left forelimb M1 at approximately 0.4 mm anterior and 1.5 mm lateral of bregma. The craniotomy was cleaned with SurgiFoam (Ethicon) soaked in phosphate-buffered solution (PBS). If the dura remained intact, a durotomy was performed with a needle.

A retrograde virus carrying a tdTomato construct (AAVretro-tdTomato, diluted to 1 × 10^13^ particles per mL in PBS, Addgene stock 59462-AAVrg) was injected in S1-fl (mice 1-3; 0.0 mm A, 2.65 mm L relative to Bregma) or the center of the forelimb representation of M1 (mice 4-6; 0.4 mm A, 1.5 mm L). 100 nL of virus was pressure injected (NanoJect III, Drummond Scientific) at a depth of 300 µm from the cortical surface over 5 min. The injection pipette was kept in place for another 3-6 minutes to ensure viral dispersion before being removed slowly over 2-3 minutes. This injection retrogradely labeled cell bodies in other areas that sent projections to the injected area. In this paper the labeling was used solely for stabilizing the imaging plane (see below) throughout the cortex.

The craniotomy was sealed with a 5 mm #1 round cover glass (Thomas Scientific) glued in place first with tissue glue (VetBond, 3M) and then with cyanoacrylate glue (Krazy Glue) mixed with black dental acrylic powder (Ortho Jet; Lang Dental). In all mice two layers of MetaBond (Parkell) were applied, and then a custom water jet-cut stainless steel head bar was affixed to the skull with black dental acrylic. Animals were awoken by administering atipamezole via intraperitoneal injection and allowed to recover at least 3 days before water restriction.

### Behavior and training

The behavioral task (Figure 2A) was a variant of the multi-target water-reaching task of Galiñanes and Huber (2023). Water-restricted mice were head-fixed with their forepaws on paw rests (bent partially threaded screws) and the hindpaws and body supported by a custom 3D-printed translucent seat. Trials were triggered after the animal held both paw rests for 100 ms. After an additional 100 ms hold period a water spout (22 gauge, modified 90-degree bent, 1-in blunt dispensing needle, McMaster) was moved into one of 15 positions in front of the mouse’s face via a set of three motors (LSM050B-T4A, Zaber). The spout was customized, with ~2 mm of the side facing away from the animal ground down; this caused the water to cling to the last few mm of the spout and not just hang from the tip. The animals were then required to hold the paw rests continuously for an additional 400-600 ms delay period to trigger the delivery of a water droplet on the spout. If the animals released the paw rests briefly during this time the delay was reset. If the paw rest release exceeded 1 second contiguously the trial was aborted and a 10 second penalty period occurred. Brief releases occurred at variable rates across animals (20-85% of trials) and delay-aborted trials rarely occurred (<1% of trials). Animals were allowed a maximum of 2000 ms to achieve paw rest contact for the entire delay; if this did not occur the spout was removed to the neutral location and a 5000 ms time-out was triggered (<3% of all trials across animals). Once the paw rests were held continuously for the variable delay time, a 4000 Hz tone was played by stereo speakers as the Go cue and a 2-3-μl droplet of water appeared on the spout. The cue tone lasted 500 ms or until the mouse made contact with the water spout, whichever came first. The mouse could grab the water droplet and bring it to its mouth to drink anytime after the tone began. The spout was removed 4000 ms after first contact, triggering a variable ITI of 3000-7000 ms, and automatically wiped against a small sponge to remove any remaining water. Both the paw rests and spouts were wired with capacitive touch sensors (Teensy 3.2, PJRC). Control software was custom-written in MATLAB R2018a using PsychToolbox 3.0.14. Touch event monitoring and task control were performed at 60 Hz using the Teensy.

The mice were trained to make all reaches with the right paw and to keep the left paw on the paw rest during reaching. For the first 1-3 days, mice were head-fixed and provided water directly to their mouths to acclimate them to head-fixation and reward collection. During this period, the animals were encouraged to make consistent contact with both paw rests to trigger water delivery to their mouths. After acclimation and consistent trial initiation, the spout was withdrawn to a position on the right side of their face intended to elicit forelimb reaching with the right arm. If animals were reluctant to reach, small droplets of water were placed on their right whisker pad with a blunt syringe to induce grooming and water collection. After animals reliably performed reaches toward the rightward spout location (80-90% success), its position was moved medially and anteriorly in gradual increments within a single session until it aligned with the midline. Once mice could reliably reach to the central spout position, two new target locations were introduced on either side (approximately 1 mm from center). Once in this three-target training stage, each target was cued in blocks of 50 trials. The size of the blocks was reduced as the mice progressed in their training until targets were interleaved randomly. After 80-90% success at the three target configuration, two additional targets were added further lateral on the left and right sides, creating a five target semicircle. The targets were then progressively spaced apart along the semicircle until they reached their final azimuthal spacings of ~1.72 target-to-target and ~6 mm total. Variation in the Z position was introduced by adding an additional row of targets above and below the existing row and gradually increasing the inter-row distance until the final Z spacing of 1 mm was achieved. This training procedure took 10-20 days. Data presented here were collected after 3-6 weeks’ experience with the task.

### Two-photon calcium imaging

Calcium imaging procedures were similar to those described previously (Grier et al. 2026). Briefly, imaging was performed with a Neurolabware 2p microscope running Scanbox 4.1 and a pulsed Ti:sapphire laser (Vision II, Coherent). Depth stability of the imaging plane was maintained using a custom plugin that made automatic movements of the objective up to every ~10s based on comparing red-channel data with an initially-acquired image stack. Offline, images were run through Suite2p to perform motion correction, region-of-interest (ROI) detection, and fluorescence extraction. ROIs were manually curated using the Suite2p GUI. Fluorescence was neuropil-corrected using a coefficient optimized by minimizing the mutual information between the corrected trace and the neuropil trace (Allen 2025). As previously described (Grier et al. 2026; Zhu et al. 2022), fluorescence was then detrended and passed through OASIS using the ‘thresholded’ method, AR1 event model, and limiting the tau parameter to be between 300 and 800 ms. To put events on a more useful scaling, for each ROI, we found the distribution of event sizes, smoothed the distribution (ksdensity in MATLAB, with an Epanechnikov kernel and log transform), found the peak of the smoothed distribution, and divided all event sizes by this value. This rescales the peak of the event size distribution to have a value of unity.

As previously described, when time-locking the data for visualization, modulation testing, or further analyses, the deconvolved events were resampled into 10-ms bins. This resampling was performed by assigning a fraction of each event into the new 10-ms bins proportionally to how much the 10-ms bins overlapped the original ~32.2 ms frames, and taking into account the exact time each ROI centroid was sampled based on its position within the field of view (FOV).

All cells were imaged between 140 and 210 µm below the cortical surface, in superficial layer 2/3. We chose this depth range both because it yielded large numbers of modulated neurons, and because GCaMP expression in these animals was lower in deeper layer 2/3 of S1. Imaging in this limited depth range therefore produced more comparable data across areas.

### Measuring modulation

To determine whether a neuron was modulated in the task, we used an adapted ZETA method (Montijn et al. 2021) as described in (Grier et al. 2026). Briefly, trial-averaged responses were compared to shuffled trial-averages computed by circularly permuting each trial by a random offset. Modulation was computed as the maximum difference between the real and shuffled data. This procedure was performed separately for each target and for locking data to the cue (−300 ms to +300 ms), lift (−200 ms to +400 ms), and first contact (−300 ms to 300 ms). For each locking event (cue, lift, and first contact), neurons with any p-values ≤ 0.01 were considered modulated. For all analyses, only neurons modulated by the relevant locking event were included. Note that this measure looks for modulation over time to any target; it is indifferent to whether the neuron exhibits tuning across targets.

### Mapping cell positions onto a common coordinate frame (Allen atlas)

Each 2P FOV was aligned based on the vasculature to an image of the entire window captured via widefield imaging. S1-fl was identified by paw vibration (Alonso et al. 2023), with imaging and preprocessing performed exactly as described in (Grier et al. 2026) except that the camera was upgraded to a pco.edge 5.5 CLHS sCMOS camera (Excelitas). The vibration map was aligned to the widefield vasculature image via shared vasculature landmarks. This composite image was then scaled and translated to best align the S1-fl activation shape to its CCF boundary, while aligning the central sinus vasculature to the midline overlapping bregma. See Figure 1-figure supplement 1 for vibration map to Allen CCF alignment across animals. 2P images from the cortical surface from each session were then scaled to 720 µm width by 980 µm height and aligned to the registered vasculature via rigid transformations. This series of transformations produced global coordinates for each ROI from its location within the 2P FOV.

### Summarizing subregions from prior literature

To produce the map in Figure 1A, outlines of subregions described by prior literature were extracted. Our review of the literature is non-exhaustive and serves simply to illustrate the heterogeneity of function that has been described in previous studies. Corresponding figures were overlaid with the Allen CCF and rigidly transformed to qualitatively match anatomical boundaries as closely as possible. Outlines were then extracted by tracing the subregion boundaries by hand.

From anterior to posterior we describe the acronyms and the corresponding citations as follows: Secondary motor cortex (M2), or MOs in the Allen CCF. Anterior lateral motor area (ALM), as originally described in Komiyama et al. (2010) and further delineated in Guo et al. (Guo et al. 2015) and Chen et al. (2017). Rostral Forelimb Area (RFA) as described in Tennant et al. (2011) and further delineated in Carmona et al. (2024). Rostral Forelimb Orofacial area (RFO) as described in An et al. (An et al. 2025), and referred to as tjM1 in Mayrhofer et al. (2019). Secondary vibrissal motor area (M2-vb) as described in Mayrhofer et al 2019 as the frontal region activated by C2 vibrissal stimulation, and called medial motor area or MM in Chen et al. (2017). Primary motor cortex (M1) or MOp in Allen CCF. Primary forelimb motor cortex (M1-fl) as described in Munoz-Casteneda et al. (2021) via tracing and characterized in Grier et al (2026). Primary hindlimb motor area (M1-hl) as traced from hindlimb musculature by Maurer et al. (2023), and sensory cortical areas by Zingg et al. (2014). Primary forelimb somatosensory area (S1-fl), or SSp-ul in Allen CCF. Primary proprioceptive and tactile forelimb somatosensory areas (S1-pro and S1-tac) as originally identified in Alonso et al. (2023). Primary hindlimb somatosensory area (S1-hl), or SSp-ll in Allen CCF. Primary trunk somatosensory area (S1-tr), or SSp-tr in Allen CCF. Primary mouth somatosensory area (S1-m), or SSp-m in Allen CCF. Primary nose somatosensory area (S1-n), or SSp-n in Allen CCF. Primary vibrissal somatosensory area (S1-bfd) or SSp-bfd in Allen CCF. Anterior visual cortex (A) or VISa in Allen CCF. Retrosplenial cortex (RSC), or RSPagl, RSPd and RSPv in Allen CCF. Cortical cervical spinal cord (CSC) and lateral rostral medulla (latRM) projections as traced in Yang et al. (2023).

### Computing map images from single-cell properties

To quantify and visualize the spatial distribution of properties of imaged neurons, we used a binning procedure to prevent biases due to uneven sampling. We binned the neurons into 80 µm-wide square pixels, then averaged the property of interest over all neurons within the pixel. Pixels that contained zero neurons were excluded from further analysis. The images were then Gaussian smoothed (s.d. 150 µm). To Gaussian-smooth the images while excluding empty pixels and accommodating irregular map edges, the final normalization for each pixel was the sum of the weights of contributing pixels, excluding empty pixels. For modulation images, each neuron was assigned a zero if not statistically modulated or a one if modulated. For PETH feature maps, each neuron was assigned its PETH feature value. The one-dimensional pixel values were then mapped onto a perceptually-linear colormap (Kovesi 2015).

### PETH feature computations

#### Response duration

We quantified the response duration of single neurons by analyzing the autocorrelation structure of their responses over time. For each neuron, we first computed the autocorrelation function *r*(*τ*) of the trial-averaged response over time *x*(*t*) to each target separately:

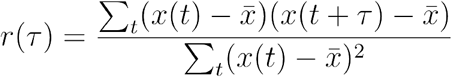

where *τ* represents time lag, and 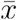 is the mean of the target-specific response over time. Only targets for which the neuron was statistically significantly modulated were included. From this autocorrelation, we determined the full width at half maximum (FWHM). The mean FWHM was calculated across all the targets for which the neuron was modulated (up to 15), resulting in a scalar measure of response duration for each neuron. Higher values of mean FWHM indicate neuronal responses that are more sustained (with a broader peak), whereas lower values indicate a more transient response.

#### Peak time variation

Some neurons exhibited firing rate peaks that were very similarly timed for all targets, while other neurons did not. To quantify this, each neuron’s PETH was first computed as described above, then for each condition (target) *c*, the peak response time 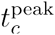 was identified as the time bin at which the trial-averaged trace reached its maximum amplitude. To prevent the inclusion of conditions with minimal modulation, we restricted this analysis to conditions whose peak amplitudes were at least 25% of the maximum peak amplitude across all 15 conditions for each neuron. The variability in peak timing was then quantified as the standard deviation of the selected peak times:

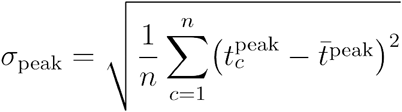

where *n* is the number of conditions meeting our inclusion criterion and 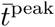 is the mean peak time across conditions above the threshold. A higher *σ*_peak_ indicates greater variability in the timing of peak responses across targets. To measure the signal-to-noise ratio for each neuron, which we used to characterize how strongly the peak time variation metric related to response consistency, we used a simple proxy from previous work (Kaufman et al. 2016). Specifically, for each neuron, we took the maximum firing rate for any time and condition minus the minimum firing rate for any time and condition, and divided this difference by the maximum s.e.m. for any time and condition.

#### Tuning sharpness

Tuning sharpness measures how selective each neuron is to some targets over others. For each neuron, we obtained the peak response *v*(*c*) for each condition (i.e., target) *c*. The tuning sharpness then computes the average response as a fraction of the max response, and subtracts this from one

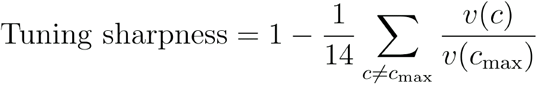

This metric ranges from near one, meaning a neuron is highly selective for a single condition, to near zero, indicating a neuron achieves a similar peak firing rate for all targets.

#### Target tuning linearity

To assess how linearly related a neuron’s response profile was to the target locations in physical space, we performed an analysis related to Representational Similarity Analysis (RSA; Kriegeskorte et al. (2008)) but limited it to linearity with respect to an ‘anchor’ target. To do so, we first computed a matrix containing the ordinal distances between all pairs of targets, and a matrix of the ‘distances’ between the firing rates for each pair of target responses (averaged over the time window from 200 ms prior to the cue to 400 ms post-cue). We then computed the coefficient of determination (R^2^) between row *m* of the target distance matrix and the corresponding row *m* of the neural distance matrix, for all rows *m*. Finally, we took as the target tuning linearity the highest of the 15 R^2^ values. Intuitively, this looks for a linear relationship between firing rate and target distance with respect to an ‘anchor’ target that maximizes the correlation. Values near one indicate a neuron whose firing scaled perfectly linearly with target location assessed in this way.

#### Tuning persistence

For each neuron, we first computed the PETH as described above. Then, for each neuron *i*, we identified the target for which the neuron achieved the greatest peak response, then found the time point *t*_*rmax*_ of that peak. An anchor tuning vector **v**_*i*_(*t*_*rmax*_) was computed, whose entries are the PETH amplitudes across targets at time *t*_*rmax*_. At every time point *t*, we measured the similarity of the instantaneous tuning profile vector **v**_*i*_(*t*) to the anchor vector by the Pearson correlation

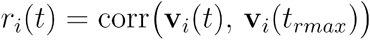

This yields a “tuning-distance” curve *r*_*i*_(*t*) that falls off as the neuron’s PETH tuning profile deviates from that at *t* = *t*_*rmax*_. To summarize these correlation plots into a single value, we computed the area under the tuning-distance curve

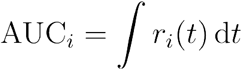

using trapezoidal integration over the sampled timepoints. Neurons with slowly decaying correlations (large AUC) achieved higher tuning persistence, whereas rapidly decaying neurons (small AUC) received low tuning persistence.

#### Local dimensionality

We estimated the effective dimensionality of local population activity by computing a common and simple metric, the participation ratio (PR), in the spatial neighborhood of each recorded neuron. This metric is limited to only considering the singular value spectrum of the local activity, but when applied exclusively to modulated neurons it provides at least a coarse idea of the variety of responses. For each neuron, we identified its *K* = 30 nearest neighbors (including the anchor neuron) by Euclidean distance in anatomical (*x, y*) coordinates. For each neighbor, the trial-averaged PETH was vectorized; stacking these vectors formed a data matrix *X* whose columns correspond to neurons. We then computed the eigenvalues λof the covariance matrix of *X*. Finally, the participation ratio for that neighborhood was taken as:

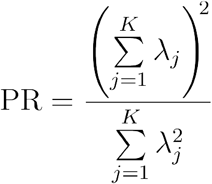

PR increases as variance is distributed more evenly across orthogonal components (higher local dimensionality), approaching 1 when variability is dominated by a single component (neuron) and approaching *K* (number of neighbors) when contributions are comparable across all components. The procedure was repeated with each neuron serving as the anchor, yielding one PR value per neuron.

### Smoothing feature distributions

In Figures 4A, 5, 8-figure supplement 1 and 9G, distributions of features were smoothed with Kernel Density Estimation (KDE). Specifically, for a distribution, the density 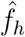 at point *x* was computed as

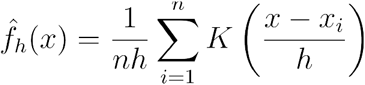

where {*x*_1_,*x*_2_,*x*_3_, …, *x*_*n*_} represent the distribution of interest, *K* is the Gaussian kernel, and the bandwidth *h* was set by Silverman’s rule (Läuter 1988):

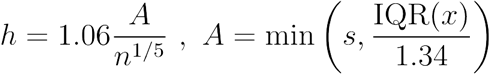

with *n* being the number of observations *s* and being the sample standard deviation.

### PCA-based alternative to feature-based analyses

As an alternative to our feature extraction method for summarizing population structure in trial-averaged activity, we applied PCA to the vectorized PETHs of all modulated neurons (the matrix of shape N x CT) and used the first 20 coefficients of each neuron as a feature space. For increased interpretability, we performed a VARIMAX rotation on the first 20 coefficients to sparsify them without distorting the space. The rotated scores’ spatial structure was visualized in Figure 5-figure supplement 1. The likelihood maps for clustering based on the PC coefficients were shown in Figure 9-figure supplement 3C-G.

### Gradient calculation

In Figure 6, we asked whether the single neuron PETH feature values changed more over the same spatial distance in some locations than others. This corresponds to a spatial derivative of the metric field. To compute this for each map in Figure 6, we first smoothed the spatial distribution of the feature map using the irregular Gaussian smooth function as described in the section “Computing map images from single-cell properties**”**. We then gridded the smoothed map values into square bins as described above, and filled missing interior values with linear interpolations of their neighbors. At each pixel, we then computed the 2D gradient magnitude using MATLAB’s centered finite-difference method. Specifically, if *M* (*x,y*) denotes the gridded feature map, then the gradient magnitude |∇*M*(*x,y*)| was calculated as:

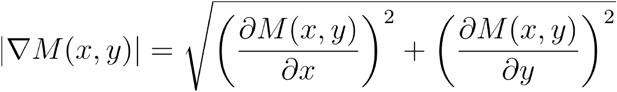

This produced a scalar map of local changes in the corresponding feature map. For the version in Figure 9-figure supplement 5, the two partial derivatives (*x* and *y*) were computed for each of the four subpopulation likelihood maps. The quadratic mean was then calculated as above over the set of individual feature (or likelihood) gradient magnitude maps.

### Random boundary analysis

In Figure 6G,H we generated random boundaries through rejection sampling. Potential midpoint coordinates were randomly selected uniformly within the bounding box of the map, and an orientation was selected from a uniform distribution of 0-360°. We then computed the endpoints for a 1 mm line segment with that center and orientation. This boundary candidate was accepted if two criteria were met. First, both endpoints must have been within a non-empty pixel. Second, at least 200 neurons must have been in the valid zones on each side of the boundary (between 50 µm and 500 µm from the boundary). To account for non-uniform distributions of neurons, we downsampled neurons on the denser side of each random boundary to match the count on the sparser side.

To assess how well we could classify neurons as above or below a random boundary using the neurons’ feature vectors, we trained a binary SVM with a radial basis function kernel and parameter optimization via Bayesian hyperparameter search. Decoding performance was evaluated using 5-fold cross-validation.

### Nonlinear dimensionality reduction

We used t-SNE to visualize the structure of our neural population’s feature space. Each neuron was represented by a five-dimensional feature vector comprising the PETH feature values described above. To improve the representation of large-scale structure, we seeded the t-SNE optimization with the top two principal components of the feature matrix (Kobak and Linderman 2021). The affinity metric was chosen to be the Pearson correlation. t-SNE perplexity hyperparameter was chosen according to the *N*/100 rule of thumb (Kobak and Berens 2019). The t-SNE space produced a two-dimensional projection of our neural data with coordinates of *Y*_*i*_ = (*y*_*i*,1_,*y*_*i*,2_) for each neuron *i*.

To produce contours in the t-SNE space for each area, we first selected the points for a given area and performed two-dimensional Gaussian KDE on these points to produce an estimate of probability density. We then drew contours (*contour* in Matlab) at the 90th, 95th, and 98th percentiles of the peak. To verify that our results were not specific to t-SNE, we also applied two-dimensional Uniform Manifold Approximation and Projection (UMAP; McInnes et al. (2020)) and obtained qualitatively similar results.

### Gaussian Mixture Modeling and cluster selection

For the analyses in Figures 8, 9 and 10, we modeled the PETH feature vectors from each area using Gaussian Mixture Models (GMMs). For each candidate number of clusters *k*, we fit a GMM with full covariance matrices using the Expectation-Maximization algorithm (fitgmdist in Matlab). To select the optimal number of mixture components, we used the Integrated Completed Likelihood (ICL) criterion (Bertoletti et al. 2015), computed as:

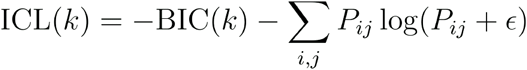

where *P*_*ij*_ denotes the posterior probability that neuron *i* belongs to cluster *j*, BIC represents the Bayesian Information Criterion (Schwarz 1978), and *ϵ* was chosen as 10^−4^. For each area, we fitted GMMs with one to ten components, and selected the fit that produced the maximal ICL.

### Likelihood computations

Following model selection for each region, we computed the log likelihood of every neuron according to the GMM of each anatomical region. The log-likelihood ℒ_*i*_ of neuron *i* given the full model for a single area was defined as:

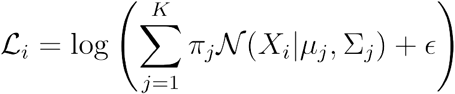

where *K* is the number of components in the GMM and *π*_*j*_ is the mixing weight for component j. In addition to the full-mixture model, log-likelihoods were calculated for each component within an anatomical area’s full mixture model to quantify the spatial footprint of each Gaussian component and to compare this spatial map across components of different anatomical areas. These log likelihoods provide a computational approach for understanding the structure of single neuron feature properties across the sensorimotor cortex.

### GMM component distances and clustering

In Figure 9B, we computed the distance between each pair of GMM components found in different areas. To do so, we computed pairwise Bhattacharyya distances between the Gaussians. This distance captures differences in the means (as a Mahalanobis-like distance), and differences in the variances and covariances (differences in the orientations, shapes, and sizes). Each Gaussian *k* was represented by its mean vector ***μ***_*k*_ ∈ ℝ^*N*^ and covariance matrix ∑_*k*_ ∈ ℝ^*N*×*N*^, where *N* is the feature dimensionality (5). The Bhattacharyya distance *D*_*B*_ between multivariate Gaussians *i* and *j* was defined as

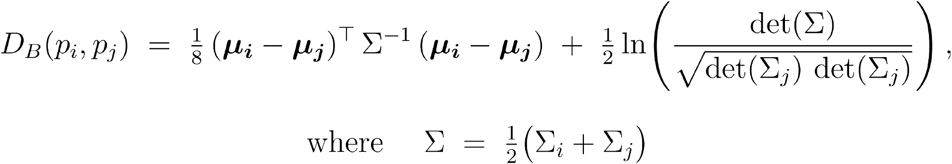

For display, the distance matrix produced by the Bhattacharyya distance was ordered using hierarchical clustering with Ward’s method (Figure 9B). The number of clusters (‘subpopulations’) was determined by simple examination of this matrix. Each cluster comprised GMM components with heavily overlapping distributions in the feature space. Clusters were further validated by comparing the spatial distributions (in cortex) of the member neurons (Figure 10E), which were very similar for the components grouped in each cluster.

As the simplest way to fairly represent the components that were grouped to form subpopulations, for further analysis we modeled each subpopulation as arising from a GMM comprising the Gaussian components that were grouped together by clustering, weighted equally.

## Acknowledgements

The authors thank P. Ravishankar for animal care assistance and L. Pinto for helpful comments on the manuscript. This work was funded by The University of Chicago, NIH-NINDS R01 NS121535 (MK), the Simons Foundation (MK), the Whitehall Foundation (MK), the NSF-Simons National Institute for Theory and Mathematics in Biology via grants NSF DMS-2235451 and Simons Foundation MP-TMPS-00005320 (MK), and NIH T32 NS121763 (HG).

## Data availability

Data will be made available on FigShare upon publication.

### Code availability

Code used to create the figures is available on GitHub: https://github.com/kaufmanlab/SGK25-public

## Supplementary figure captions

**Figure 1-figure supplement 1. Widefield window images and vibration maps aligned to the Allen CCF for all mice**. Shown as in Figure 1B.

**Figure 2-figure supplement 1. Reach-to-grasp-to-drink centroid trajectories show moderate target-specificity**. (A) Single-trial finger centroid trajectories locked to lift onset as in Figure 2C, shown from a posterior to anterior perspective, one session per mouse. Target positions not shown. Single-trials highlighted for one target for mouse 3, the same mouse shown in Figure 2C. (B) Each trace quantifies how different reaches were on average. For each possible pair of trials, we computed the Euclidean distance between finger centroid XYZ positions at each time point. Red traces show the average distance across trials to different targets and black traces show the average distance across trials to the same target. Shading shows SEMs across pairwise distances. In many cases the shading is occluded by the mean trace. Horizontal bars above the traces show time points with significantly different distributions of within vs. between distances at p < 0.05 from a two sample Kolmogorov-Smirnov test.

**Figure 3-figure supplement 1. Modulation maps for all task events**. (A) Binned and smoothed modulation map as in Figure 3B, but for cells modulated by Cue (left), Lift (center) or First-contact (right). Cue alignment used neural data from 200 ms before to 400 ms after cue. Lift alignment used neural data from 200 ms before to 400 ms after lift. First-contact alignment used neural data from 300 ms before to 300 ms after first spout contact. (B) Binary matrix indicating which area-area comparisons had significantly different proportions of cue-modulated cells. Black squares indicate rejection of the null hypothesis at p < 0.005 using a two proportion z-test.

**Figure 5-figure supplement 1. Principal component scores after VARIMAX rotation for each neuron**. Plotted as feature maps using the same method as Figure 5.

**Figure 8-figure supplement 1. Feature value histograms for all GMM components and areas**.

**Figure 9-figure supplement 1. Highest likelihood PETHs from each subpopulation**. Two PETHs are shown from each subpopulation, for a total of four per area. The two PETHs from each subpopulation for each area had the highest likelihood values across cells from that area. PETHs were chosen in a principled way as the highest likelihood neurons for each subpopulation within each area, with a max firing rate of at least 33 events/s. This approach identified the only two Anterior cells in S1-hl and zero cells in S1-tr; thus the S1-tr entries were left blank.

**Figure 9-figure supplement 2: Separation of subpopulation distributions in feature space**. (A) Feature vectors for all neurons within a subpopulation were projected onto the axis that best linearly separated the distributions, for each pairwise comparison between subpopulations. The axis was obtained from a logistic regression and the cross-validated performance of the model is shown for each comparison. Note that subpopulation membership was determined by GMM classification, which is nonlinear, while logistic regression is linear. Because of both this model mismatch and cross-validation of the logistic regression, classification performance on the projection axis is less than 100%. (B) t-SNE plot of feature vectors as in Figure 7 but colored by subpopulation membership. (C) Time to half-max distributions for each subpopulation, plotted as in Figure 4A.

**Figure 9-figure supplement 3. Additional subpopulation validation analyses**. (A) t-SNE as in Figure 7C but using the top 20 PCs of the PETHs as inputs instead of feature vectors. Multimodality was again strongly present. (B) Left, subsampled population of neurons chosen from an overlap zone in M1, to analyze for discrete subpopulations while avoiding spatial somatotopy. *Left*, t-SNE embedding of this M1 population using the PETH feature space. *Right*, t-SNE embedding of the same M1 population using the PCs. Both methods yielded clear multimodality. (C-G) Same as Figure 8B-F but using top 20 PCs instead of PETH features.

**Figure 9-figure supplement 4. GMM fit to all PETH feature vectors together, agnostic to anatomical areas**. One GMM was fit to the feature vectors of cells from all 5 main areas. Each map plots the likelihood for all cells to each of the three components of this area-agnostic GMM.

**Figure 9-figure supplement 5. Gradients of subpopulation likelihood maps**. (A-D) Gradients as in Figure 6 but calculated using the subpopulation likelihood maps depicted in Figure 9C-F instead of the PETH feature maps. As in Figure 6, each subpopulation had a distinct gradient with boundaries approximately aligned with borders between anatomical regions, and other boundaries approximately separating somatotopic representations. (E) The quadratic mean of the four subpopulation gradient maps. This pooled gradient map closely resembles the pooled map in Figure 6F.

