## Supplemental figures for "Neural activity profiles reveal overlapping, intermingled subpopulations spanning area borders in mouse sensorimotor cortex"

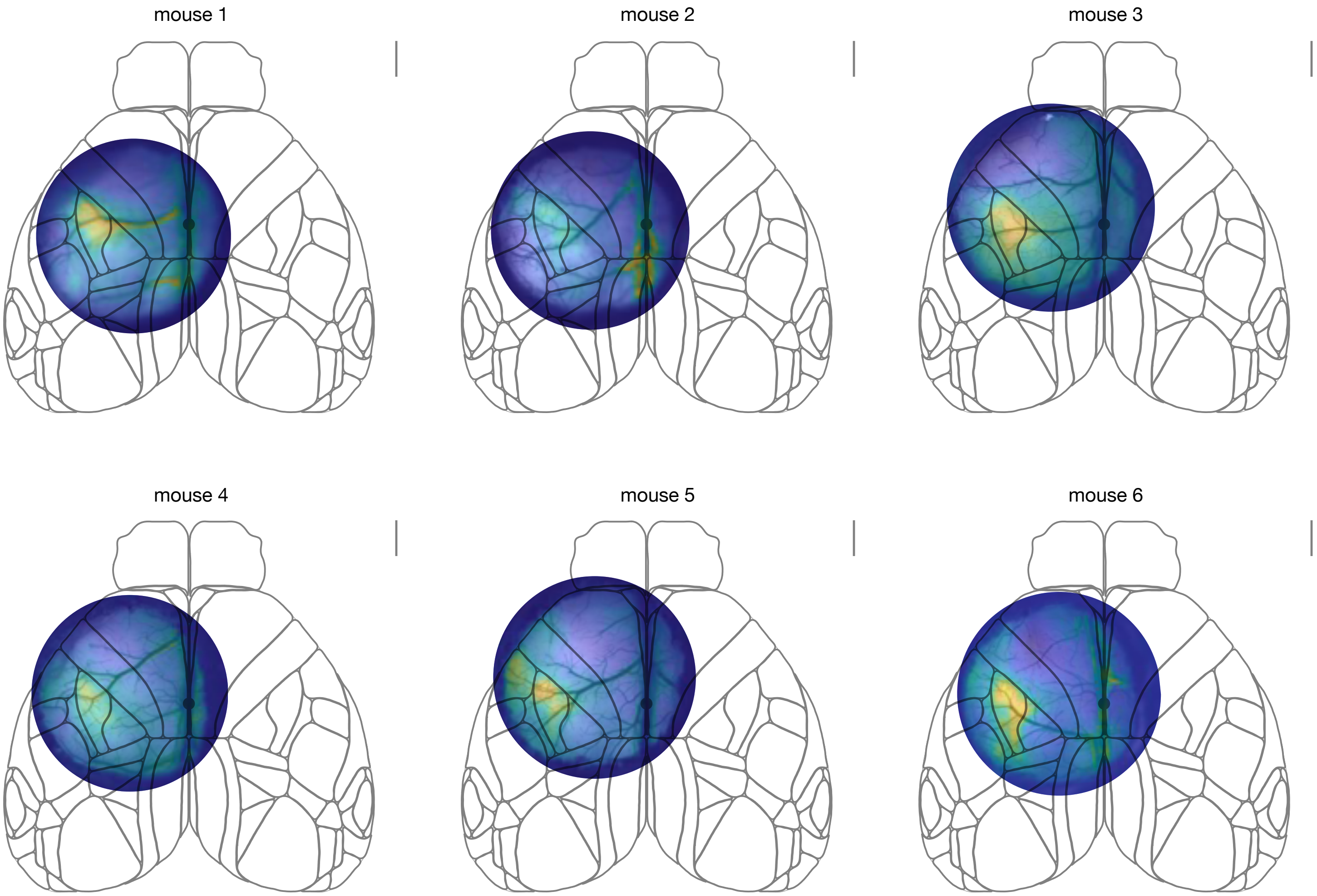


**Figure 1-figure supplement 1**

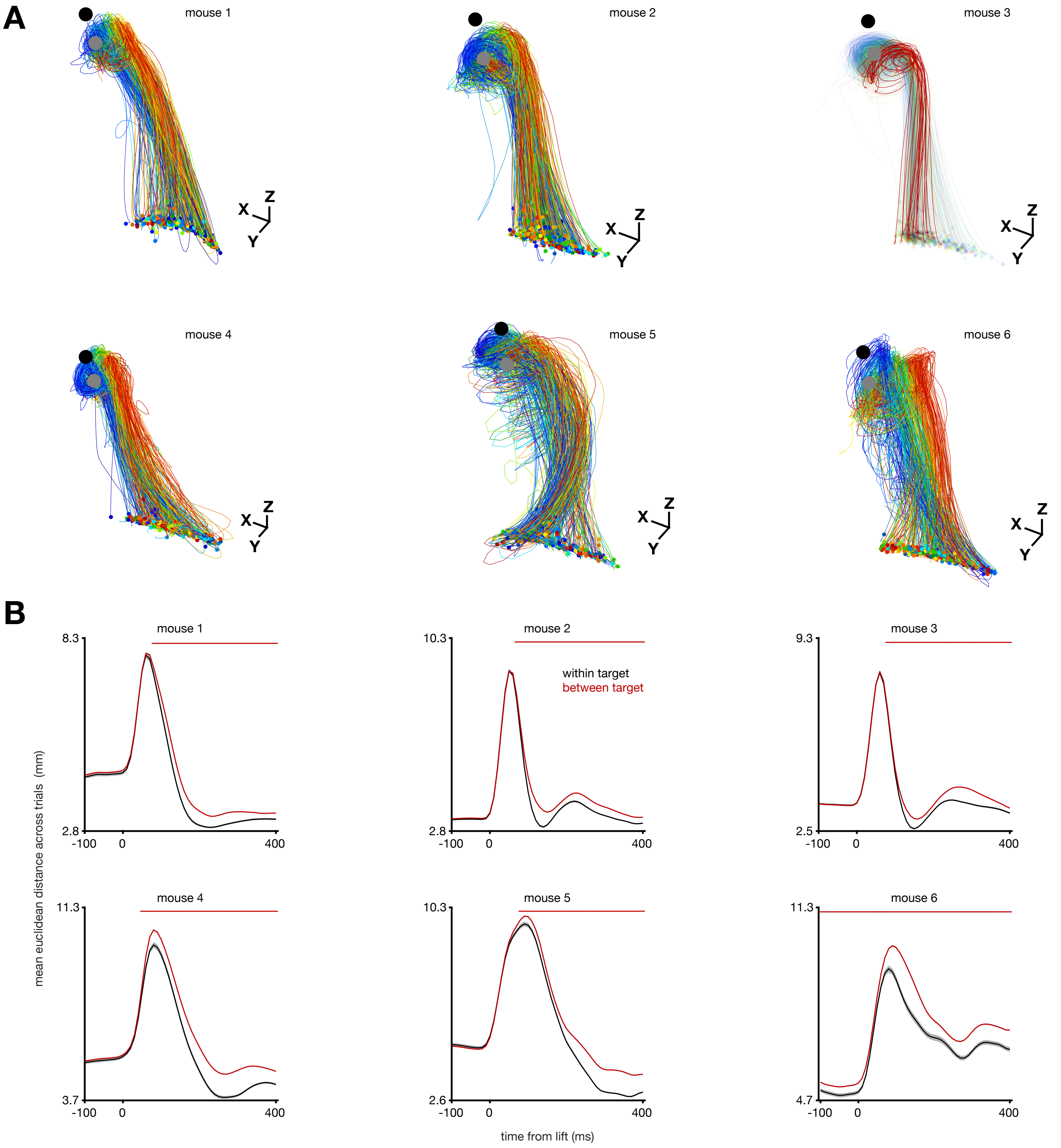


**Figure 2-figure supplement 1**

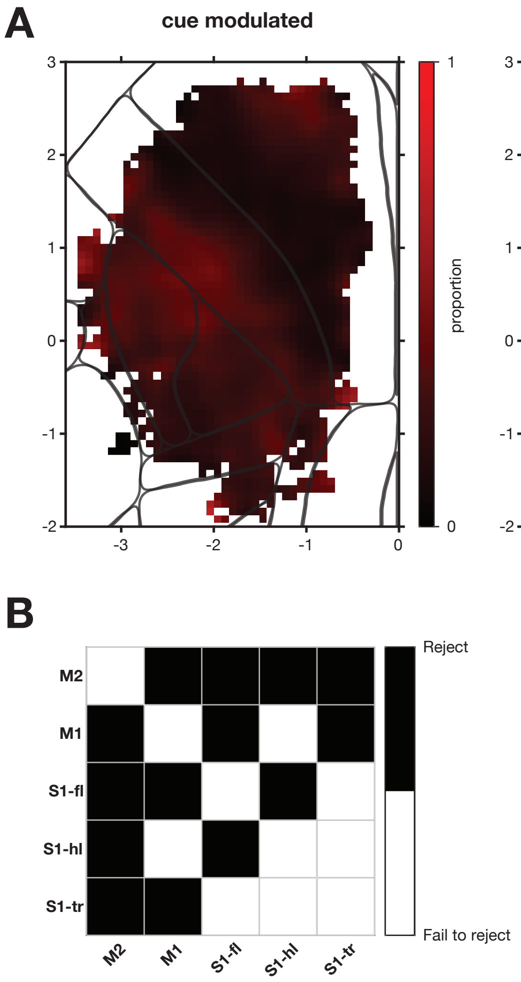


**Figure 3-figure supplement 1**

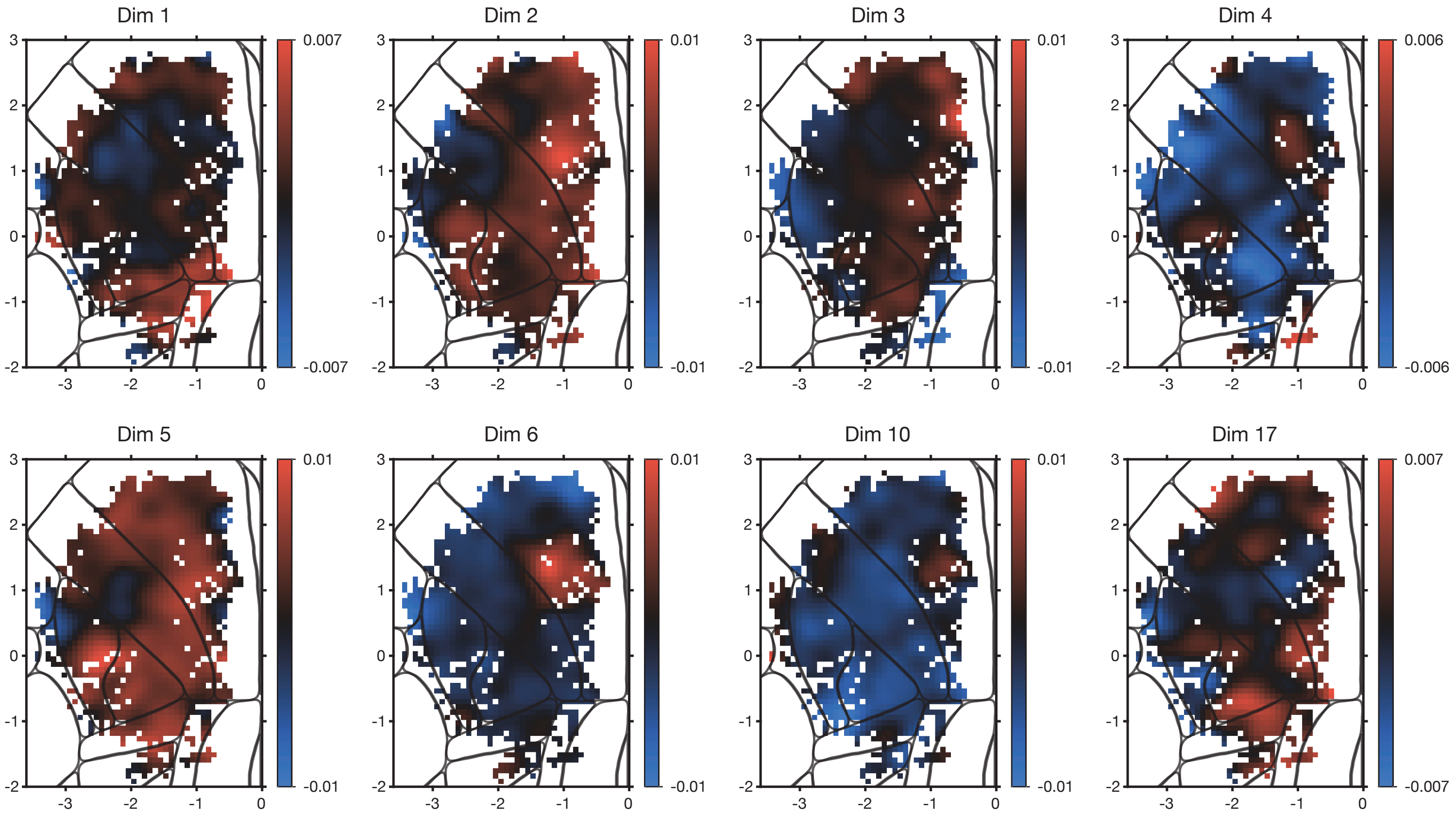


**Figure 5-figure supplement 1**

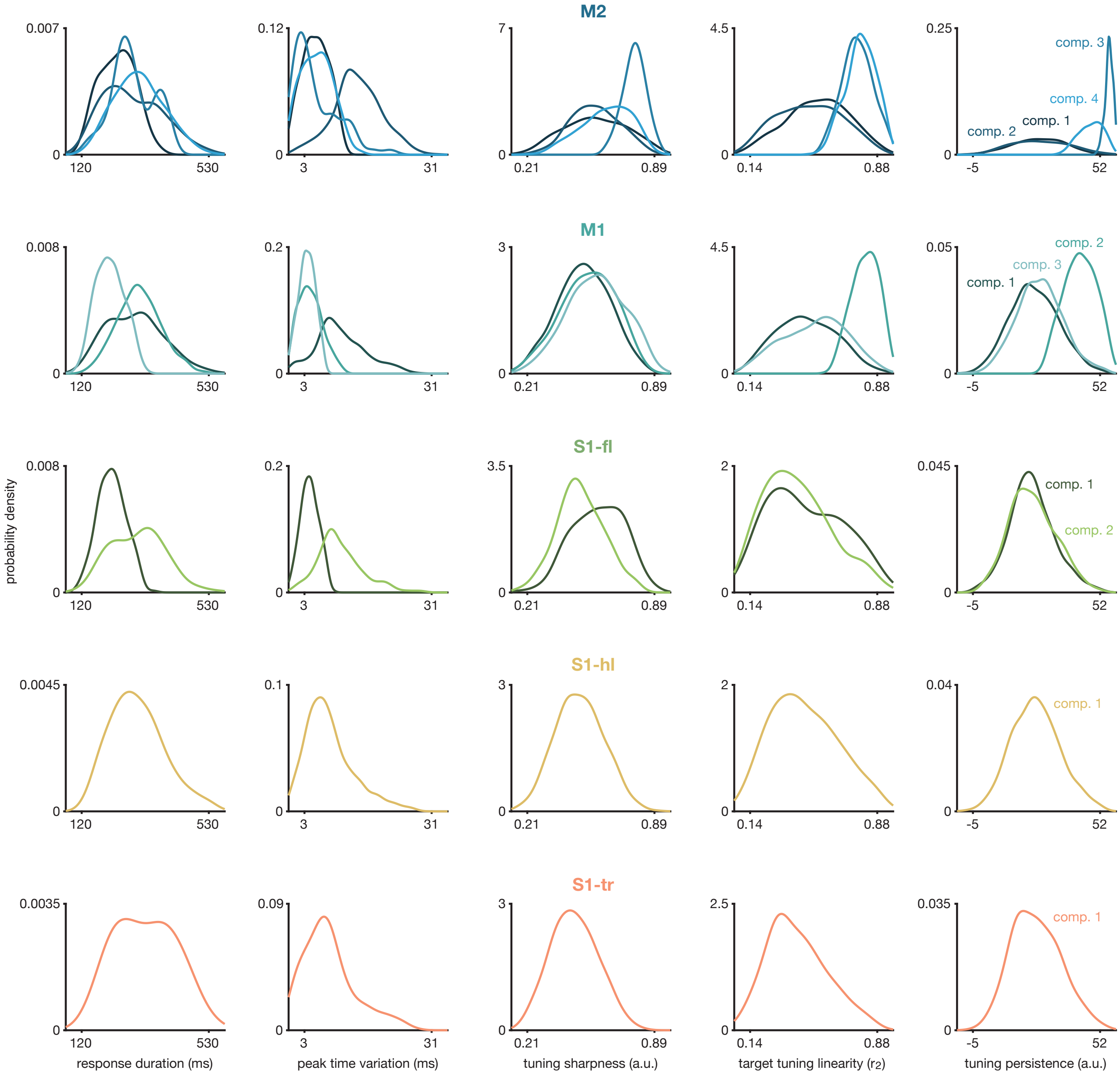


### Figure 8-figure supplement 1

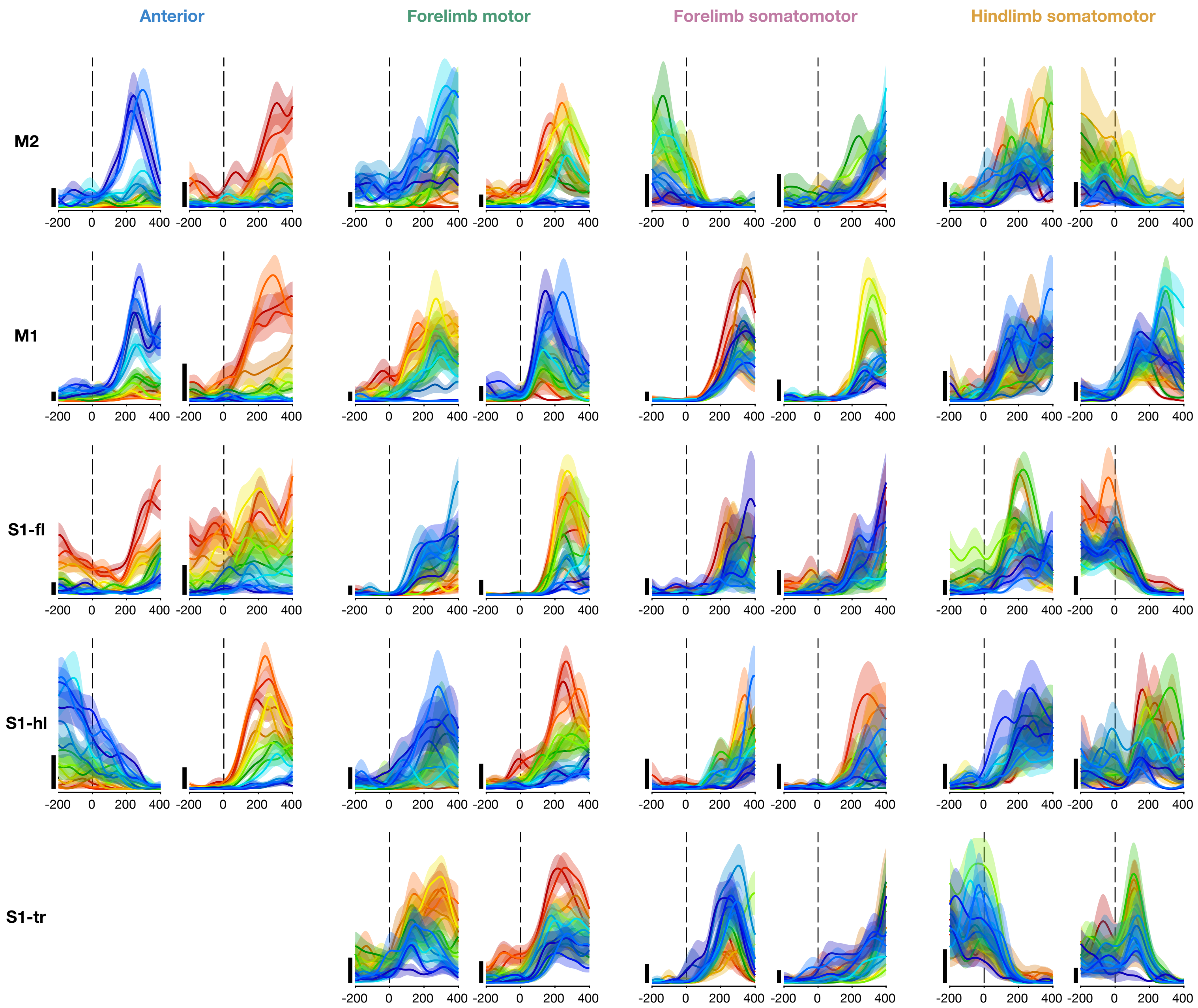


**Figure 9-figure supplement 1**

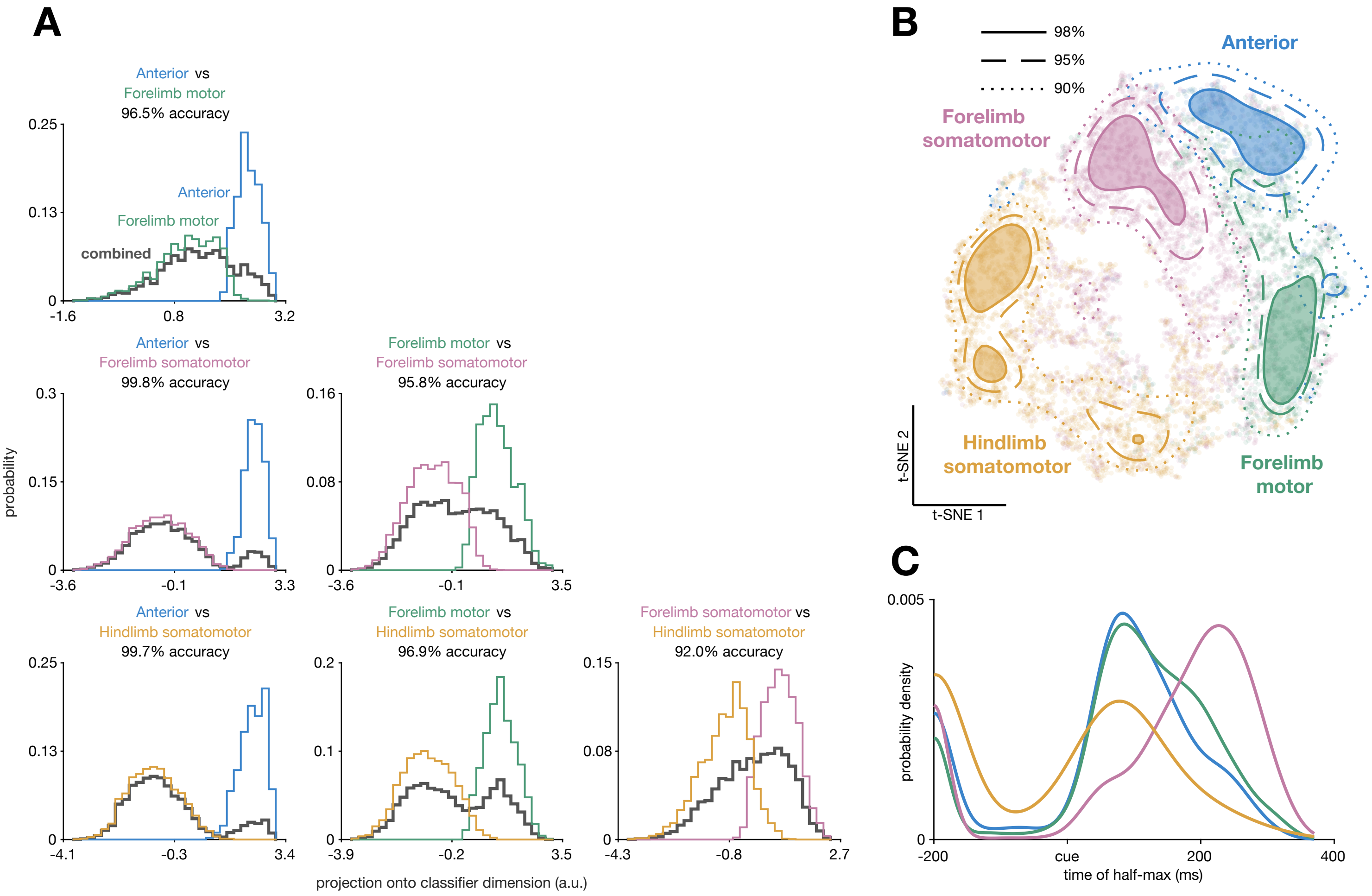


**Figure 9-figure supplement 2**

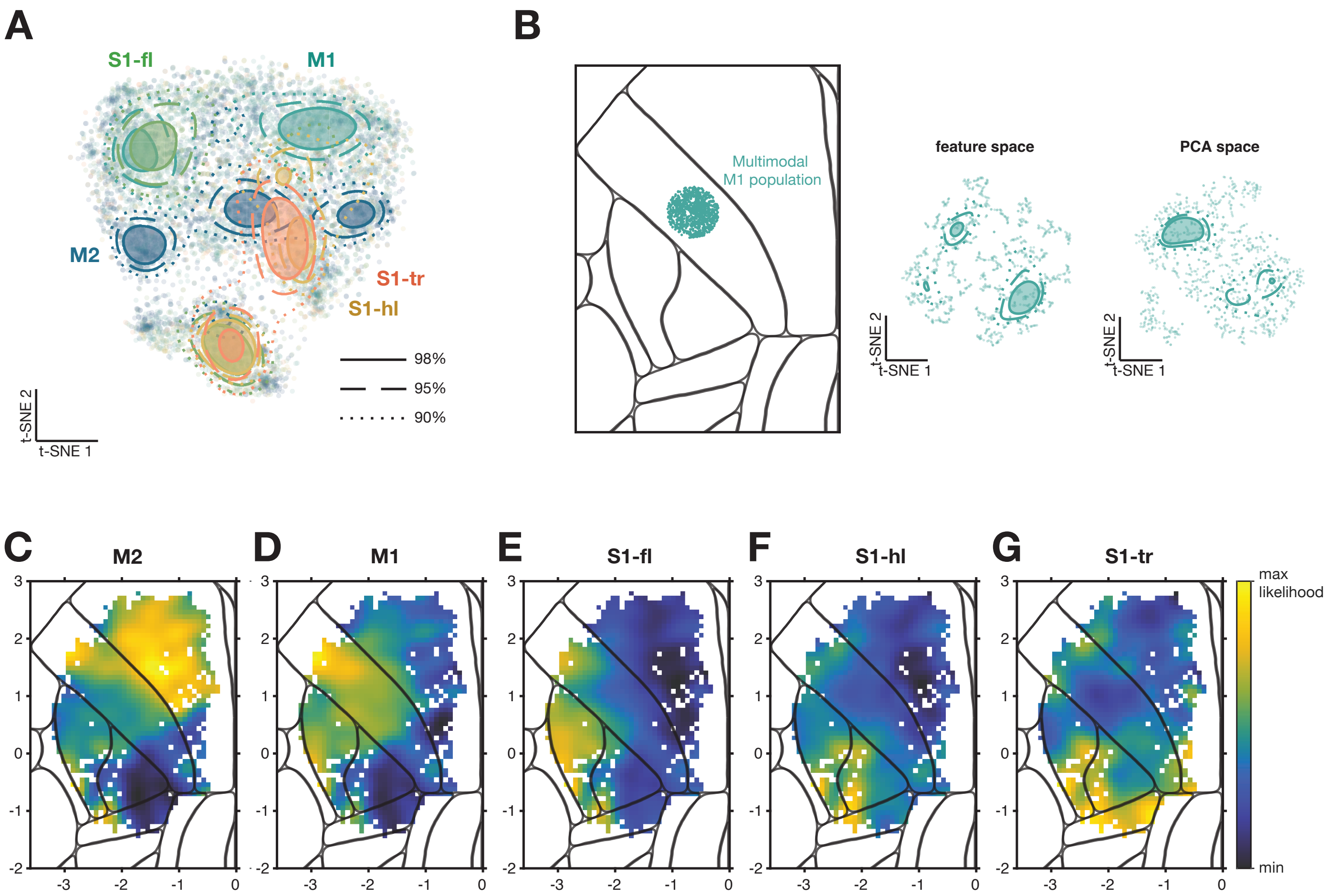


**Figure 9-figure supplement 3**

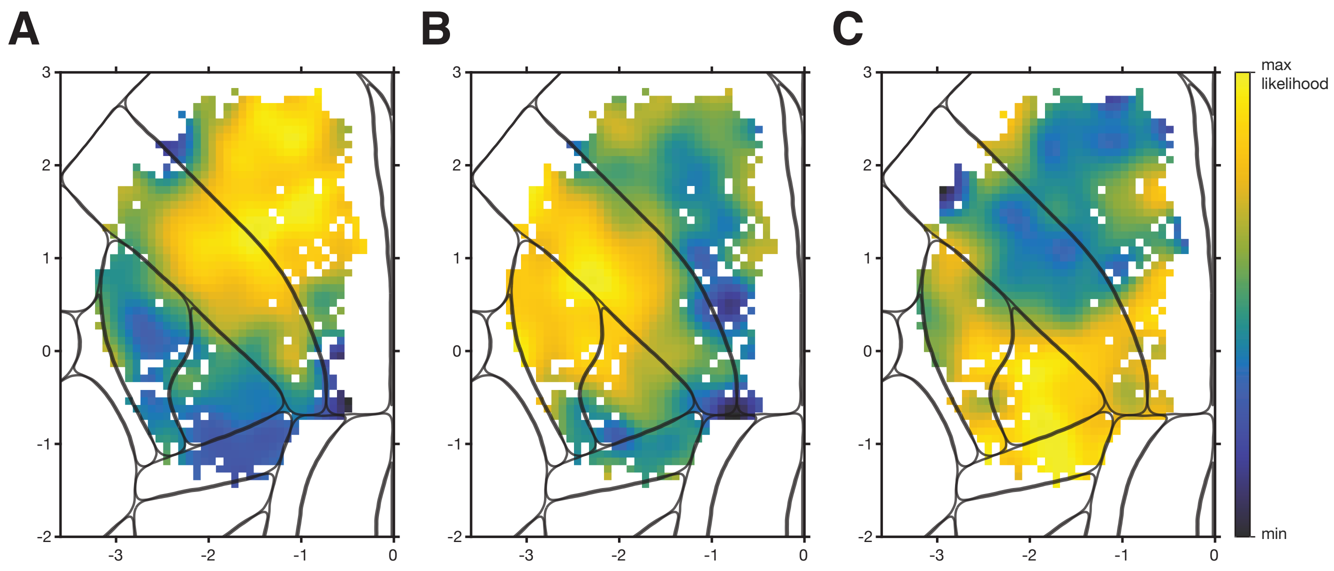


**Figure 9-figure supplement 4**

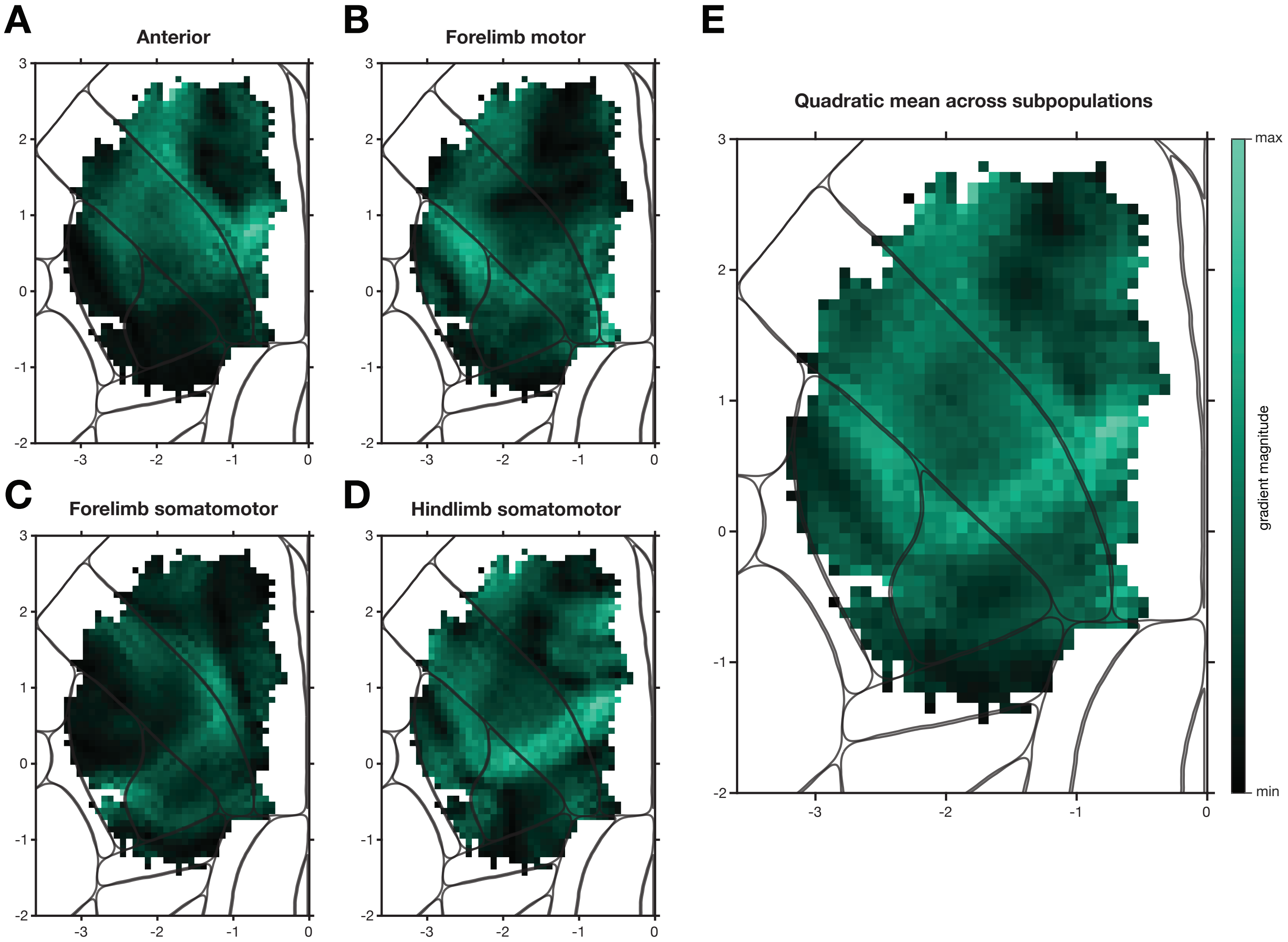


**Figure 9-figure supplement 5**
